# CURTAIN – A Unique Web-based tool for exploration and sharing of MS-based proteomics data

**DOI:** 10.1101/2023.07.25.550405

**Authors:** Toan K. Phung, Kerryn Berndsen, Tran Le Cong Huyen Bao Phan, Miratul M. K. Muqit, Dario R. Alessi, Raja S. Nirujogi

## Abstract

To facilitate analysis and sharing of mass spectrometry (MS)-based proteomics data we created tools called CURTAIN (https://curtain.proteo.info) and CURTAIN-PTM (https://curtainptm.proteo.info). These enable the non-MS expert to interactively peruse volcano plots; deconvolute primary experimental data to individual replicates that can be visualized in bar charts or violin plots allowing statistical analysis; and export of plots in SVG format. They also permit assessment of experimental quality by correlation matrix and profile plot. Within CURTAIN, the user can analyze domain structure, AlphaFold predicted structure, reported interactors, relative expression, disease and pharmaceutical links, and mutagenesis information on all selected hits. Moreover, CURTAIN-PTM permits the comparison of all identified PTM sites on protein(s) of interest with PTM information contained within selected databases. For phosphorylation site analysis CURTAIN-PTM links with the kinase library to predict upstream kinases that phosphorylate sites of interest. We provide examples of the utility of CURTAIN and CURTAIN-PTM in analyzing how targeted degradation of the PPM1H Rab phosphatase that counteracts the Parkinson’s LRRK2 kinase impacts cellular protein levels and phosphorylation sites. We reanalyzed a ubiquitylation dataset, characterizing the PINK1-Parkin pathway activation in primary neurons, revealing new data of interest not highlighted previously. CURTAIN and CURTAIN-PTM are free to use and open-source and will enable researchers to share and maximize the analysis and impact of their proteomics data. We advocate that differential expression proteomic data should be published containing a shareable CURTAIN web-link, allowing readers to better explore their data.

**Significance Statement:** To enable non-experts to better share and explore mass spectrometry data, we have generated using open-source software, interactive tools termed CURTAIN and CURTAIN-PTM. These tools enable users’ to save their analysis sessions with a sharable unique web-link, enabling other researchers to visualize and further analyze these datasets. These links can also be reported in publications allowing readers to further survey the reported data. We discuss benefits for the research community of publishing proteomic data containing a shareable web-link.

## Introduction

Mass spectrometry (MS)-based proteomics is the go to method to analyze proteomes and posttranslational modifications and is transforming all areas of biological investigation (1). Recent advances in sample preparation methodologies include nano-liquid chromatography and ultra-sensitive MS instrumentation that enable the identification and quantification of >10,000 protein groups in a single shot injection, with amounts as little as 200 ng or less of cell or tissue extracts (2, 3). MS-based proteomics has hugely impacted our understanding of signal transduction pathways by enabling accurate pinpointing and quantification of global patterns of post-translational modifications including protein phosphorylation (4, 5), glycosylation (6), acetylation (7) and ubiquitylation (8). Recently developed trapped ion-mobility coupled time-of-flight MS platforms make it possible to undertake ultra-sensitive proteomics even at a single cell level (9). Most MS data is now being acquired in a data dependent (DDA) or data-independent acquisition (DIA) mode generating half a million MS and MS/MS events that take up to 10 GB per file (10).

Differential expression proteomic MS-based data is commonly presented in a static volcano plot format (11). These are routinely generated by an MS expert following processing of the primary MS (raw) data from all replicates and experimental conditions employing a sophisticated suite of analysis software including MaxQuant (12), MS-Fragger (13), Spectronaut (14) or DIA-NN (15). The output from these programmes are complex text files that are subsequently analyzed in statistical analysis programs such as Perseus (16) or MSstats (17). This analysis requires significant expertise and involves the use of scripts and packages from Python and R.

The output from this analysis that is routinely shared or submitted to journals for publication are complex spreadsheets containing thousands of lines and dozens of columns together with non-interactive visualizations such as volcano plots. These volcano plots frequently contain a vast amount of useful information, and it is usually only possible for the presenter to highlight a small subset of the data that is of particular interest to them. Much data that could be of interest to other researchers is not highlighted nor straight-forward to visualize. It is challenging for the non-MS expert to subsequently retrieve and reanalyze the published data contained within a volcano plot image, and analyze the sets of proteins that they are most interested in. This requires access to the algorithm and statistical analysis output files and typically such analysis needs to be performed by a suitably trained expert. These analyses also require access to sophisticated software packages, many which are not free to use open-source, and need to be downloaded on a high-end computer. It is also demanding for the non-expert to deconvolute the data presented on a volcano plot so that the primary data from each of the replicate experimental samples can be visualized and further analyzed.

Similar issues arise in analyzing volcano plots from post translational modification (PTM) analysis. It is frequently necessary to explore all the experimental PTM data available for a protein of interest, within each dataset. It is also important to visualize the location of each identified PTMs on the protein(s) of interest and how these modifications are impacted by experimental conditions. It is also necessary to be able to easily compare experimental PTM data with publicly available PTM databases such as PhosphoSitePlus (18), Protein Lysine Modification Database (PLMD) (19), CarbonylDB (20), GlyConnect(21) and Uniprot (https://www.uniprot.org/) (22). Finally, it is useful to analyze the amino acid sequence motifs encompassing each identified PTM.

To address the above issues, we have generated, two open-source software, interactive tools termed CURTAIN (https://curtain.proteo.info) and CURTAIN-PTM (https://curtainptm.proteo.info). CURTAIN and CURTAIN-PTM necessitate no local computer server installation and operate through the internet on any computer equipped with a modern web browser. These tools are designed to provide users the option to save their analysis sessions with a sharable unique web-link, enabling other researchers to visualize and further analyze these datasets. These links can also be reported in publications allowing readers to easily explore and verify the quality of the reported data. We provide examples of the utility of CURTAIN and CURTAIN-PTM in analyzing how targeted degradation of the PPM1H Rab phosphatase that counteracts the Parkinson’s LRRK2 kinase (23, 24) and how this impacts on cellular protein levels using CURTAIN, and phosphorylation sites employing CURTAIN-PTM. To further demonstrate the utility of CURTAIN-PTM in analyzing ubiquitylation data, we have re-analyzed a previously reported data set (25), characterizing how ubiquitylation of mitochondrial proteins is regulated by the Parkin E3 ligase. This reanalysis revealed interesting data not highlighted in the previous study. We discuss benefits for the research community of publishing proteomic data containing a shareable web-link, allowing readers to better explore data with open-source tools such as CURTAIN or CURTAIN-PTM.

## Results

### CURTAIN, a web-based tool for visualization and analysis of MS-based proteomics data

To facilitate analysis and sharing of complex MS data, we have created a web-based open-source tool called CURTAIN (https://CURTAIN.proteo.info/<u>#/</u>). The detailed front and backend framework of CURTAIN is described in the Methods section and summarized in Figure 1A. It allows interactive exploration of proteomic datasets which can be shared via a web-link. CURTAIN is designed to be used by non-MS-expert end users, once the analyzed MS data is uploaded by the MS expert who undertook the experiment.

**Figure 1:**
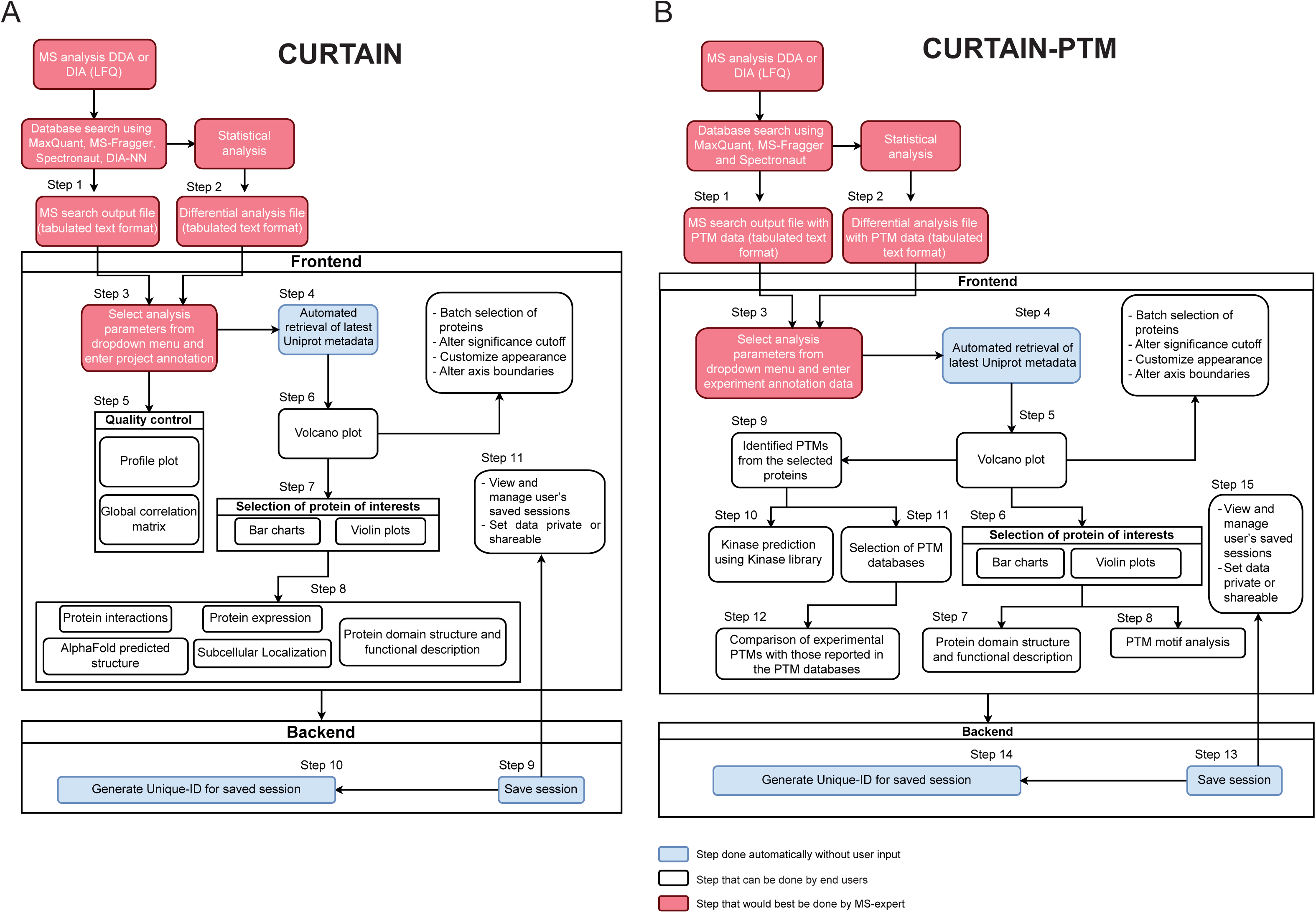
Workflow and for importing, analyzing, and exploring processed MS-based total proteomics data with CURTAIN and CURTAIN-PTM. For further details refer to the Methods and Result section.

CURTAIN requires two MS input files to be uploaded that we recommend is undertaken by the MS expert who undertook the study. First is the database search output file, in tabulated text format, of the primary MS data obtained from either MaxQuant (12), MS-Fragger (13), Spectronaut (14) or DIA-NN (15) in either data dependent acquisition (DDA) (LFQ and TMT) as well as Data Independent acquisition (DIA) (LFQ) modes. (Step 1, Fig 1A). Second is a differential analysis file also in tabulated text format, that is obtained from statistical analysis program such as Perseus (16) or MSstats (17) (Step 2, Fig 1A). In addition, the file columns to be analyzed and compared in the input file need to be selected and this is undertaken using a dropdown menu (Step 3, Fig 1A, Dataset S1, SFig 2A). At this stage the metadata associated with the experimental data can be uploaded in the same format as used by the PRIDE database including project title, project description, sample processing protocol, data processing protocol, MS Instrumentation details, name and affiliations of researchers involved in the project (26) (Step 3, Fig 1A, SFig 2B). On opening a file, CURTAIN automatically retrieves the latest Uniprot metadata for each protein within the uploaded data (Step 4, Fig 1A). At this stage we recommend that the MS expert sends a weblink to end users to enable them to explore and analyze the data. All the subsequent steps can be readily performed by the non-specialist end user.

**Figure 2:**
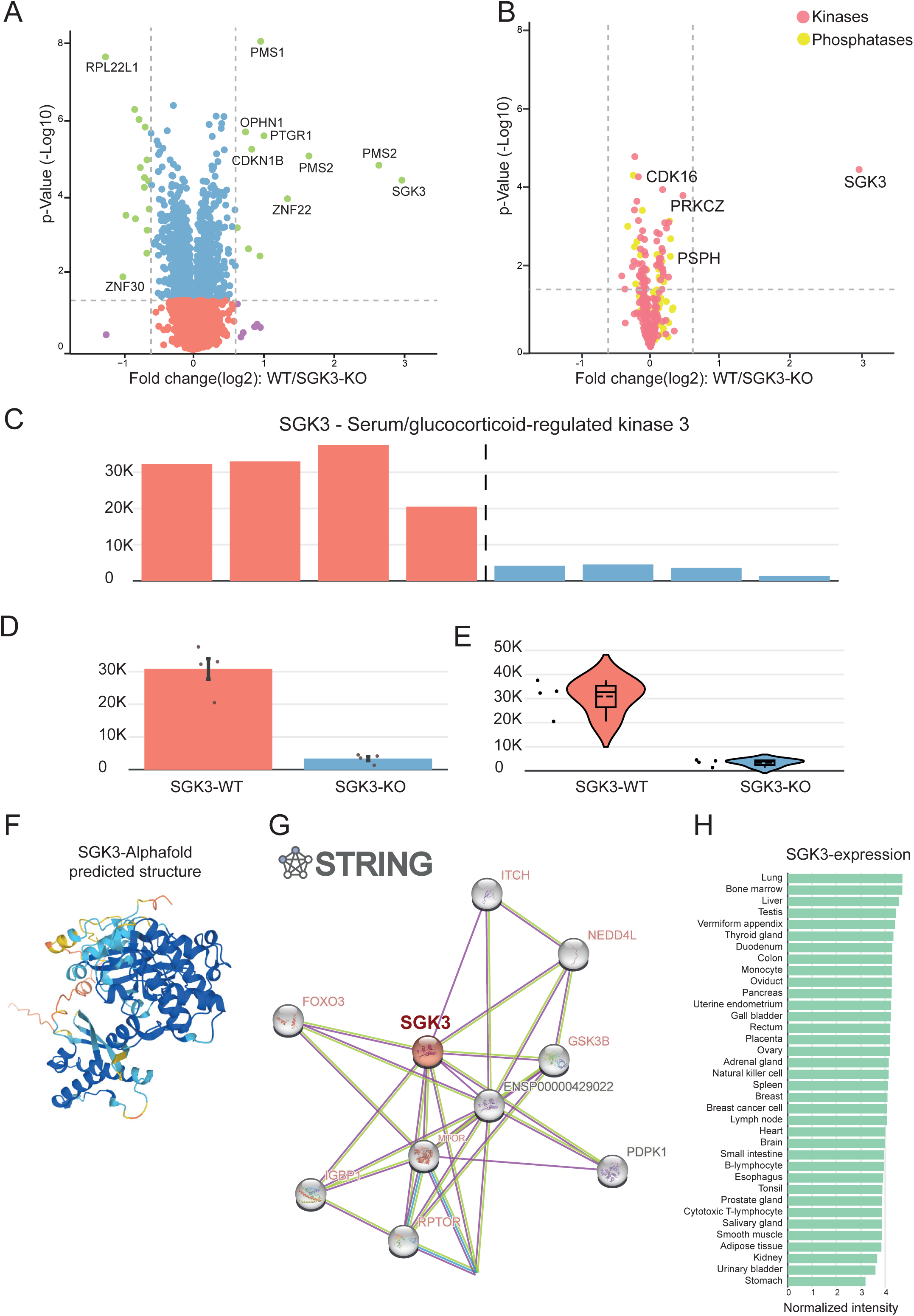
Functionalities of CURTAIN tool. A) Example of an interactive CURTAIN volcano plot employing previously reported proteomic data comparing wild type (n = 4) and SGK3 (n = 4) knock-out HEK293 cells (PRIDE data set identifier: PXD014561) (43). This data can be perused using the web-link https://curtain.proteo.info/#/b9e85c9e-5d26-41f2-8474-18e3985fa804. The colored filled dots denoting the significance and fold change cut-off values (Red: non-significant and no difference; Blue: significant and no difference; purple: non-significant but show increase/decrease in protein levels and green: significant and show increase/decrease in protein levels), proteins of interest can be selected and labeled and further analyzed within CURTAIN. B) Example of using CURTAIN to select sets of proteins of interest using inbuilt lists in this case kinase (pink) phosphatases (yellow) Denoting the custom selection of various in-built pathway components (Red filled dots: Kinases and Yellow filled dots: Phosphatases). All the non-selected hits have been hidden for clarity. The link to this data is https://curtain.proteo.info/#/aab22416-663a-40c3-a2e6-1839ff0d4dbd. C to H) SGK3 hit was selected for further analysis. The relative levels of SGK3 in 4 replicates of two experimental conditions (wild type and knock-out) were viewed in individual bar graphs (C) or combined average bar graph (± SEM or SD) (D) or violin plot (E) and the CURTAIN link for this data is https://curtain.proteo.info/#/4e792c03-099b-4b1b-bd4e-18afe4204f2c. (F) Reported STRING SGK3 interactors with gene names are color coded to depict the presence/absence within the experiment and increase/decrease in fold-change levels (Dark red: Detected/Increase in expression; Blue: Detected/Decrease in expression; Light red: Detected/NO-difference and Grey: Not detected in the dataset). G) Protein expression data of SGK3 derived from ProteomicsdB. F) Predicted AlphaFold structure of SGK3.

CURTAIN enables end users to assess the quality of proteomic data by generating profile plots (SFig 3A) and global correlation matrix (SFig 3B) of uploaded data (Step 5, Fig 1A). On the profile plots the relative levels of specific proteins of interest can be displayed across all samples (SFig 3A). CURTAIN displays the differential analysis data from each set of experiments as an interactive volcano plot displaying the fold-difference (X-axis, typically in Log2) and p-value significance (y-axis, typically in -log10) (Step 6, Fig 1A, Fig 2A). The end user can then customize the Volcano plot display color including color blind friendly options, labeling, annotation (including font size and color selection) and altering axis boundaries (Step 6, Fig 1A, Fig 2B). It also allows modification of fold change and significance cutoff to be represented by vertical and horizontal dotted lines respectively (Step 6, Fig 1A, Fig 2A, 2B). In the default settings hits, that pass the cutoff values, are represented in a different color but this can be modified as required.

**Figure 3:**
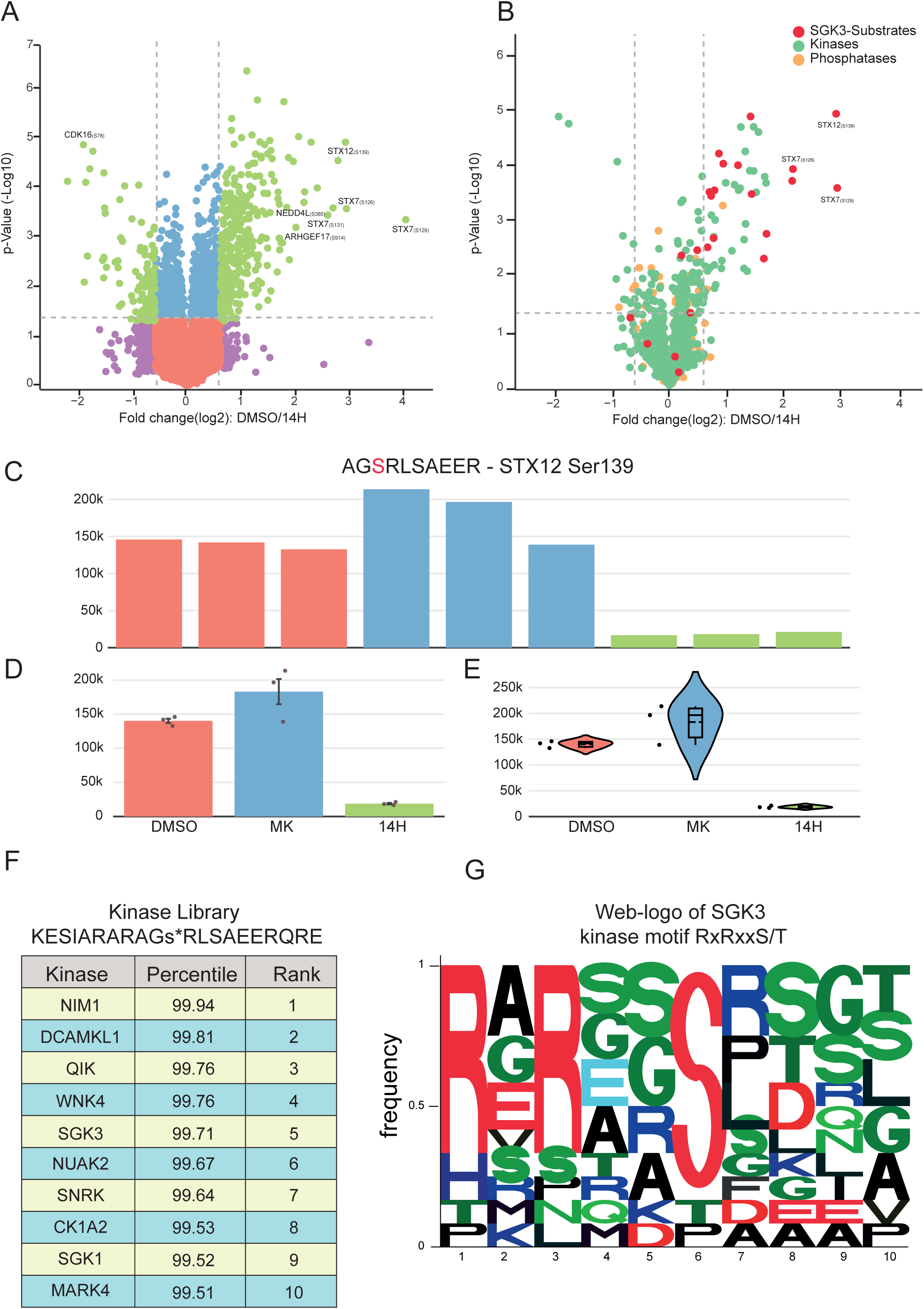
Functionalities of CURTAIN-PTM tool. A) Example of an interactive CURTAIN-PTM volcano plot using previously reported phospho-proteomic data of wild-type HEK293 cells that had been serum starved overnight and the treated with either no inhibitor (DMSO control n = 3) or a selective pan Akt isoform inhibitor (1 M MK-2206, MK abbreviation, 1h, n=3) or a pan SGK isoform inhibitor (3 µM 14H, 1 h, n = 3) prior to stimulation with IGF-1 (50 ng/ml, 15 min) in the continued presence of the inhibitor (PRIDE data set identifier: PXD014561) and data analyzed using MaxQuant (43). This data can be perused using the web-link https://curtainptm.proteo.info/#/d79cff8f-cd8f-4f4a-b082-7f7807f8a3d4. The colored filled dots denoting the significance and fold change cut-off values (Red: non-significant and no difference; Blue: significant and no difference; purple: non-significant but show increase/decrease in protein levels and green: significant and show increase/decrease in protein levels), proteins of interest can be selected and labeled and further analyzed within CURTAIN-PTM. B) Example of using CURTAIN-PTM to select sets of PTM sites, namely SGK3 substrates (red) kinase (green) and phosphatases (yellow). SGK3 substrates were selected using a custom selection of hits whilst kinases and phosphatases are from the inbuilt list. All non-selected hits have been hidden for clarity. The link to this data is https://curtainptm.proteo.info/#/a1566c36-91f3-45e2-b955-881194681ba5. C to G) The SGK3 substrate, STX12 (Ser139) was selected for further analysis. The relative levels of STX12 (Ser139) in 3 replicates of two experimental conditions (± 3 µM 14h) were viewed in individual bar graphs (C) or combined average bar graph (± SEM or SD) (D) or violin plot (E) and the CURTAIN link for this data is https://curtainptm.proteo.info/#/d79cff8f-cd8f-4f4a-b082-7f7807f8a3d4. (F & G) The sequence encompassing Ser139 site on STX12 was automatically submitted to the kinase library database to predict possible kinases that may phosphorylate this residue (31) (F) or the weblogo tool (40) (G) and the CURTAIN-PTM results output from these analyses are shown.

For more in depth analysis, the end user selects data of interest, for example a set of proteins which are changing between experimental conditions (Step 7, Fig 1A, Fig 2B). For each selected protein, CURTAIN allows visualization of the primary data for all replicates for each condition directly as a bar chart either for separate (Step 7, Fig 1A, Fig 2C) or combined bars ± SEM or SD (Fig 2D). It also allows data to be presented as a violin plot format ± SEM or SD (Step 7, Fig 1A, Fig 2E). It also allows further statistical analysis to be undertaken for example ANOVA test between experimental groups using the jstat package (https://jstat.github.io) (Step 7, Fig 1A, Fig 2E).

Another feature of CURTAIN is that it provides the option for users to learn more about the identified proteins (Step 8, Fig 1A). For all selected hits, CURTAIN allows visualization of the Uniprot functional domain structure which also provides a concise functional description of what is known about the selected protein as well as predicted/known subcellular localization (22), alphafold structure (27) (Fig 2F), known or predicted interactors derived from STRING (28) (Fig 2G). Another feature for the interactome database is that it reports interactors of selected hits that are identified within the uploaded data, highlighting proteins whose levels change between experimental conditions (Red for higher, blue for lower, pink present in dataset but levels unchanged and black not detected in dataset) (Fig 2G). CURTAIN permits searching of the Interactome Atlas (29), expression profile from ProteomicsDB (30) (Fig 2H). If of interest, CURTAIN will also link selected hits to the disease, pharmaceutical and mutagenesis data available within the Uniprot database.

CURTAIN also allows for the facile batch selection of a group of proteins of interest to the end users’. We have separated these into 4 categories namely diseases (e.g. Parkinson’s Alzhemer’s), enzymes (e.g., kinases, phosphatases), organelles (e.g. Golgi, lysosomes) and pathways (e.g. LRRK2, PINK1), that are listed in (Dataset S2) and these can readily be updated and added to. CURTAIN also allows users’ to select groups of proteins within their dataset or more generally that are of particular interest and to save these as a set to facilitate further analysis of these proteins. To further facilitate viewing of selected proteins of interest, CURTAIN permits “hiding” of all other non-selected proteins in the Volcano plot (Fig 2B).

Each analysis session on CURTAIN can also be saved and shared with a unique weblink (Step 11, Fig 1A). The person(s) receiving this link can further analyze and download the data. All graphical outputs from CURTAIN can be exported in SVG format and further modified in programs such as Adobe Illustrator. Web-links to the CURTAIN data including volcano and violin plots can be included in the figure legend of publications allowing readers to easily access, view and analyze the data further. The source code for CURTAIN can be downloaded in GIThub (see methods), hosted on a local server, and modified for additional functionalities. To help organize and keep track of different projects, CURTAIN allows users to log in using ORCID and all the different user sessions across projects are saved within the user account. Saved session data can be maintained private or shared with a weblink (Fig 1A). CURTAIN can also be used without account creation and sessions saved and retrieved using a unique weblink.

### CURTAIN-PTM, for visualization and analysis of post-translational modification proteomics data

To analyze MS-based posttranslational modification (PTMs) data, we have also developed CURTAIN-PTM (https://curtainptm.proteo.info/#/). CURTAIN-PTM deploys the same backend as CURTAIN. The search output and differential files containing PTM information are uploaded (Fig 1B). Currently CURTAIN-PTM is optimized to use output from MaxQuant PTM and search both DDA (LFQ and TMT) and DIA (LFQ) mode. It also links to MS-Fragger but requires a minimal post processing step of the data, employing a custom-made script that we have developed termed MS-Fragger to CURTAIN-PTM CONVERTER (https://doi.org/10.5281/zenodo.8138524), prior to uploading in CURTAIN-PTM, best undertaken by an MS-expert (Fig 1B). For our PTM site analysis we normally upload Class-I, high confidence sites (>0.75 probability), but it is possible to upload files containing lower probability sites if required. CURTAIN-PTM displays the differential PTM analysis data from each set of experiments as an interactive volcano plot that can be further analyzed, modified, exported, and saved as described for CURTAIN (Fig 3A). The end user selects a set of modified peptide(s) (e.g. phosphorylated or ubiquitylated) of interest, for example those whose levels are modulated between experimental conditions (Fig 3B).

CURTAIN-PTM also allows for the deconvolution and visualization of the primary data for all replicates of each condition as a bar chart with error bars (± SEM or SD) (Fig 3C, 3D) or violin plot format (Fig 3E). For phosphorylation site analysis, CURTAIN-PTM links to Kinase Library (https://kinase-library.phosphosite.org/site) (31), thus providing predictions for kinases that may phosphorylate this selected site, based on sequence analysis and knowledge of kinase substrate specificities (Fig 3F). The user can also select a group of peptides for sequence motif analysis using the weblogo tool (Fig 3G).

For each selected peptide, CURTAIN-PTM provides the gene name along with the PTM residue number(s) derived from the search algorithm (Fig 3A). CURTAIN-PTM also allows the user to link back to the protein from which it is derived from, allowing for all experimentally identified PTMs found within that protein to be visualized on a linear protein sequence and compared against selected publicly available databases (SFig 4A). The Uniprot database (22) is the default option, but other databases including PhosphoSitePlus (18) can also be selected (Dataset S3).

**Figure 4:**
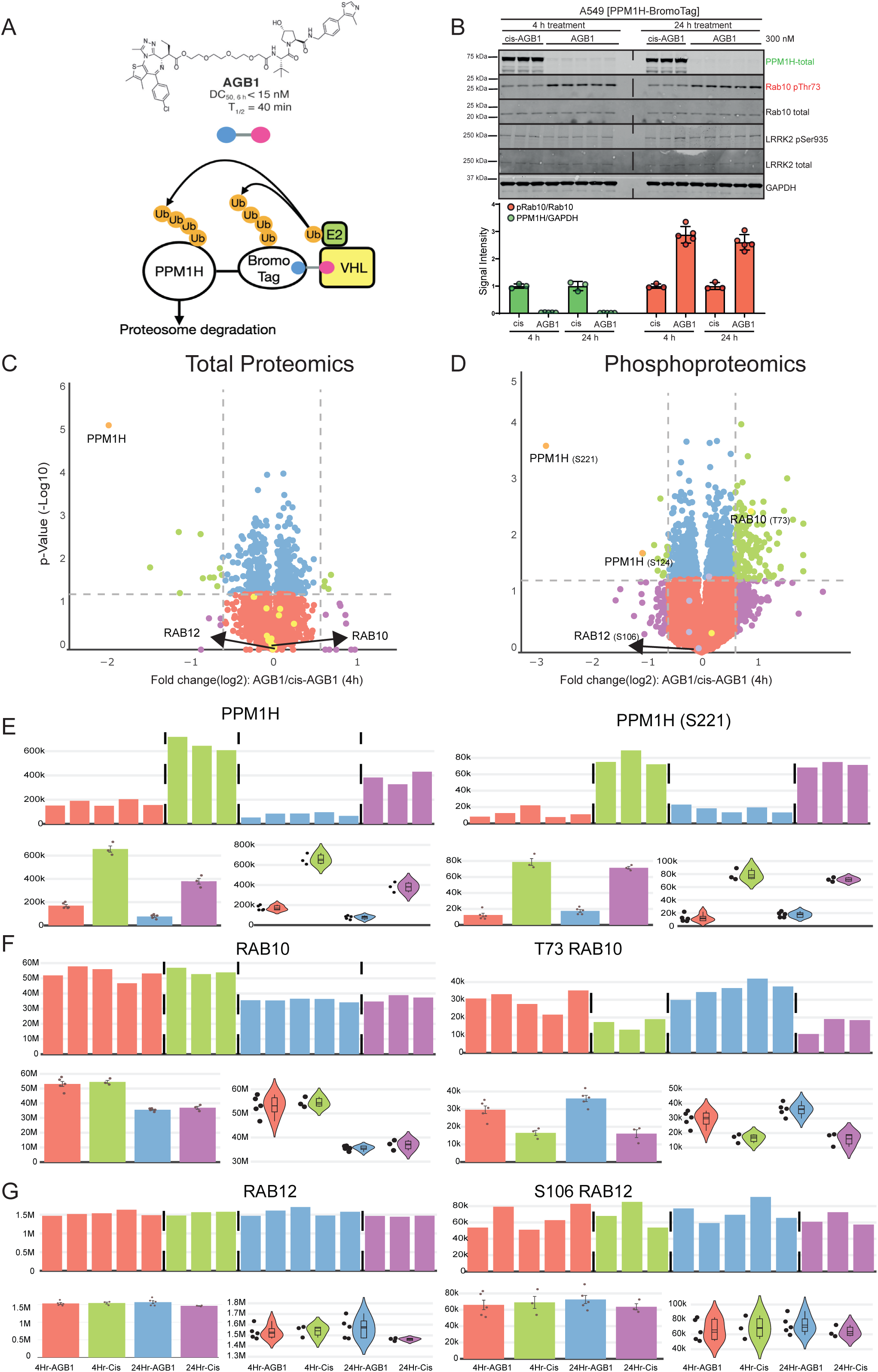
Use of CURTAIN and CURTAIN-PTM to analyze the impact of cellular depletion of the PPM1H protein phosphatase. A) Schematic depicting the mechanism by which the AGB1 compound induces the targeted degradation of the CRISPR knock-in PPM1H-BromoTag. B) Knock-in A549 PPM1H-BromoTag cells were treated with 300 nM cis-AGB1 (inactive control compound) or AGB1 for 4 h or 24 h prior to cell lysis. 20 μg whole cell lysate was subjected to quantitative immunoblot analysis with the indicated antibodies. Membranes were developed by the LICOR Odyssey CLx Western Blot imaging system. Quantitation of the ratio of phospho-Thr73/total Rab10 and PPM1H/GAPDH was obtained using Image studio software. Each lane represents lysate obtained from a different dish of cells. Individual data points of the replicates within the figure are shown, with the error bars representing the standard deviation of the mean between the replicates. C) The lysates produced from the experiments described in Fig 6B in which cells were treated for 4h with cis-AGB1 or AGB1 were subjected to total and phospho-proteomic analysis with data being analyzed with MS-Fragger and MaxQuant. CURTAIN generated volcano plot are presented for the total proteomic (C, https://curtain.proteo.info/#/f4b009f3-ac3c-470a-a68b-55fcadf68d0f) and Phospho-proteomic (D, https://curtainptm.proteo.info/#/32837924-c5bb-41a2-9d30-b5839c012243) and proteins of interest highlighted in black. The same color coding is used as described in Fig 3A. E-G) The primary total and Phosphosite intensities of indicated hits were shown as bar graphs and violin plots.

It should be noted that the residue numbering of PTMs obtained from MaxQuant and other analysis packages depends on the splice variant selected by the search algorithm, which may differ from the canonical sequence used by other databases or commonly discussed in the literature. CURTAIN-PTM lists numbering based on that used by the original search algorithm and annotates this as “experimental data” (SFig 4A). CURTAIN-PTM then performs a multiple sequence alignment between the identified PTM site and the canonical isoform listed on the selected database (SFig 4A). If there is a difference in sequence between the Uniprot and experimental sequence due to splice variant numbering, CURTAIN-PTM displays the sequence alignment and provides the selected database PTM residue number in addition to the experimental residue position (SFig 4B). When other PTM databases are selected for comparison with the experimental data, the PTM residue number derived from the selected database(s) will be displayed. Some databases such as PhosphoSitePlus enable user-selected splice variants to be analyzed. CURTAIN-PTM color codes residue positions, purple indicates that the experimental data overlaps with PTM identified in the selected database(s), red is that experimental PTM is not observed in selected database(s), blue depicts data that is present in the selected databases, but not in the experimental data, and a peptide selected for further analysis is displayed in green (SFig 4). Sites that change significantly between experimental conditions are highlighted with an asterisk (SFig 4).

### Example of use of CURTAIN and CURTAIN-PTM to analyze proteomic and phosphoproteomic analysis

To demonstrate the utility of the CURTAIN and CURTAIN-PTM tools, we employed a BromoTag targeted protein degradation strategy (32), to rapidly deplete the endogenous levels of the PPM1H phosphatase, that we had previously been shown to dephosphorylate LRRK2 phosphorylated Rab10 (23). The BromoTag was linked to the C-terminus of PPM1H using CRISPR/Cas9 gene-editing technology as the NH-terminus of this enzyme is involved in membrane binding (24) (Fig 4A). Homozygous knock-in cell lines were selected and verified by immunoblot and DNA-sequencing analysis (SFig 5A-5D). Homozygous PPM1H-BromoTag knock-in cells were treated for 4h or 24h with the AGB1 selective degrader compound or as a control with the inactive cis-AGB1 analogue (32). Immunoblot analysis confirmed that PPM1H- BromoTag levels were reduced by >95% with the AGB1 compared to the cis-AGB1 treated cells (Fig 4B). As expected, this was accompanied by an increase in pRab10 levels as judged by immunoblotting with a phospho-specific antibody (Fig 4B). As expected, the total levels of Rab10, LRRK2 and LRRK2 biomarker phosphosite pSer935 were unaffected by AGB1 treatment (Fig 4B).

**Figure 5:**
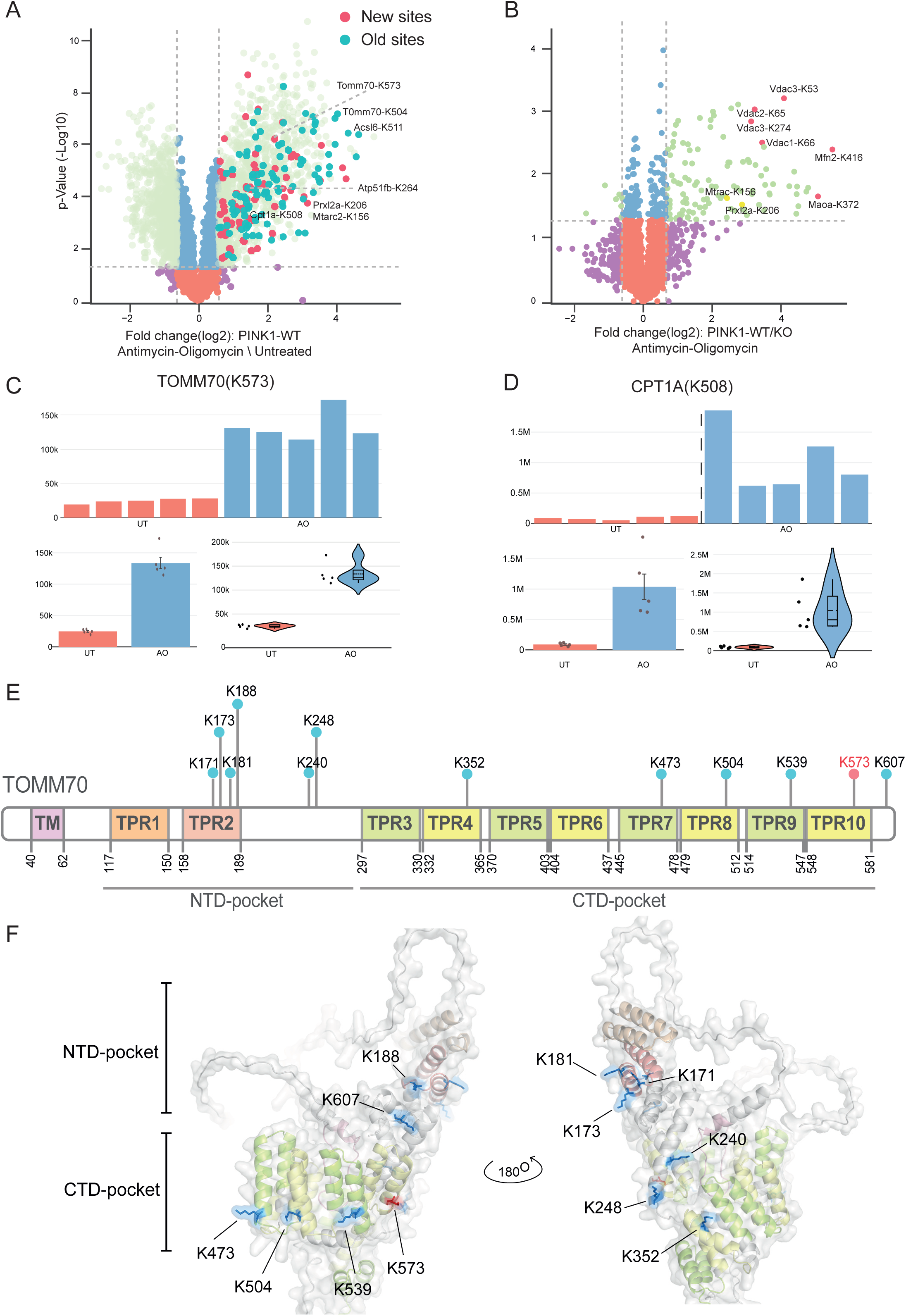
Visualization of Parkin mediated ubiquitylation in neurons using CURTAIN-PTM. The MS raw data from a previously published study (MassIVE dataset identifier MSV000087639) (25) was reanalysed using MS-Fragger search algorithm. (A) Wild type C57BL/6J primary cortical neurons were depolarized ± antimycin A (10 μM)/oligomycin (1 μM) (AO) stimulation for 5 hours or (B) wild type and PINK1 knock-out were depolarized ± AO for 5 hours. Membrane-enriched lysates were subjected to diGLY capture proteomics as described previously (25). CURTAIN-PTM volcano plots generated https://curtainptm.proteo.info/#/85970b1d-8052-4d6f-bf67-654396534d76 (A) https://curtainptm.proteo.info/#/cffbae16-abb0-418d-8244-801e7bd46bc0 (B). The color coding is described in Fig 3A, and new highlighted hits are in dark red and analyzed further in (C & D) (TOMM70-K573, CPT1A-K508) and SFig9A, 9B, whilst previously highlighted hits are shown in dark blue and analyzed in SFig9C, 9D, 9E. (E) Schematic representation of the domain arrangement of TOMM70 (M. musculus), highlighting the Trans-membrane region (TM) and TetraTricopeptide repeat (TPR) domains as annotated in UniProt. Ubiquitylated lysines (K) are indicated. (F) Structural representation of TOMM70 (M. musculus) highlighting the localization of lysine residues (previously identified ubiquitin sites in blue and novel sites in red.The structure was acquired from AlphaFold and analyzed in PyMOL (version 2.5.2). The color scheme employed corresponds to the domains depicted in the schematic representation of 9E.

We next undertook a 16-plex Tandem Mass Tags (TMT)-based quantitative proteomic and phosphoproteomic analysis as described in the Methods, of PPM1H-BromoTag A549 cells treated with AGB1 and cis-AGB1 for 4 and 24h. The MS proteomic data was searched using the MS-Fragger algorithm (13) which identified 8,609 unique proteins (Dataset S4). The phosphoproteomic data was analyzed using MaxQuant (12) which identified and quantified 31,808 phosphosites and 25,203 Class-I phosphosites (Dataset S4). Statistically significant differentially regulated proteins and phosphosites were determined using Perseus (16). The result files from these analyses were uploaded to CURTAIN and CURTAIN-PTM using the parameters described in (Dataset S1). CURTAIN generated profile plots revealed the same distribution of proteins in each sample, indicating high quality data (SFig 6A). Levels of Rab10 were similar in all samples whereas as expected, the levels of PPM1H were reduced in AGB1 compared to cis-AGB1 treated cells (SFig 6A). A CURTAIN generated global correlation matrix revealed ∼0.98 correlation between each replicate in the different experimental conditions lysed at the same time points (SFig 6B). Cells lysed at different time points displayed a reduced 0.96 correlation suggesting slight batch effects which is to be expected (SFig 6B). CURTAIN generated volcano plots for both the AGB1 4h (Fig 4C-CURTAIN, Fig 4D-CURTAIN-PTM) and 24h (SFig 7A-CURTAIN, SFig 7B-CURTAIN-PTM) time points. As expected, PPM1H was the protein most significantly reduced by AGB1, at both the 4h and 24h time points (Fig 4C and SFig 7A). Analysis of the primary PPM1H protein data confirms a marked reduction of PPM1H levels in all replicates for AGB1 treated samples (Fig 4E). A number of other proteins were moderately impacted by AGB1 and CURTAIN generated violin plots for these are shown in (SFigure 8). Batch selection of known LRRK2 pathway proteins suggests that other than PPM1H, other known pathway components were unchanged (yellow circles, Fig 4C). Based on the cutoff values used, phosphosite analysis revealed that the levels of S221 and S124 phosphosites were enhanced at the 4h (Fig 4D) and 24h (SFig 7B) time points respectively. As expected from the immunoblotting data, the phosphopeptide encompassing LRRK2 Rab10 phosphorylation site (pTh73) is enhanced ∼2-fold at both the 4h and 24h time points of AGB1 treatment (Fig 4D and 4D). LRRK2 also phosphorylates Rab12 at Ser106, but previous data suggests that PPM1H does not regulate dephosphorylation of Rab12 in vivo (23) as these proteins localize at different sites within the cell (24). Consistent with this, levels of the phosphopeptide encompassing Rab12 phosphorylated at Ser106 was not altered in PPM1H- depleated cells (Fig 4G). A violin plot of selected other phosphorylation sites that were most enhanced in the PPM1H depleted cells at both 4h and 24h time points are presented in (SFig 9). These results illustrate how CURTAIN-PTM can mine and re-use phospho-proteomics data but further work is required to establish whether these phosphosites represent direct PPM1H substrates.

### Example of use of CURTAIN-PTM from proteomic and ubiquitylation site analysis

We next deployed CURTAIN-PTM tools for re-analysis of a previously published dataset of PINK1-Parkin mediated ubiquitylation sites in mitochondria of primary C57BL/6J mouse cortical neurons (25). Tryptic peptides derived from mitochondrial enriched extracts of mouse neurons treated with or without 10 μM Antimycin A / 1μ M Oligomycin (AO) for 5 h to induce activation of Parkin followed by a diGLY immunoprecipitation-enrichment of ubiquitylated peptides which were quantified using a TMT-based approach (25) (Fig 5A). Similar re-analysis was performed for a dataset of wild-type (WT) and PINK1 knockout (KO) neurons (Fig 5B) (25).

The ubiquitylation data was re-analysed using MS-Fragger which identified and quantified 6,035 ubiquitylation sites. Statistically significant differentially regulated ubiquitylation sites were determined using Perseus. The result files from these analyses were uploaded to CURTAIN- PTM using the parameters described in (Dataset S5). We generated a CURTAIN volcano plot for AO 5 h treatment versus untreated (Fig 8A), which enabled the facile visualization of previously reported AO-induced ubiquitylated sites (light green circles, Fig 5A) including Parkin-dependent mitochondrial substrates (dark blue circles, Fig 5A). CURTAIN also generated volcano plot for AO 5 h treated PINK1 WT versus KO neurons depicting previously reported PINK1-dependent ubiquitylated sites including VDAC1 (K66) and Mfn2 (K416) (Fig 5B).

Interestingly, re-analysis elaborated ubiquitylated sites on proteins not previously reported including Atp51fb (K264), Mtarc2 (K156) and Prxl2a (K206) (red circles, Fig 5A) and of which the latter two sites are PINK1-dependent (Fig 5B). In future work it would be interesting to assess whether Mtarc2 and Prxl2a are direct Parkin substrates. Furthermore, re-analysis elaborated new sites on previously reported Parkin substrates including TOMM70 (K573) (Fig 5C) and CPT1A (K508) (Fig 5D). In addition, other new ubiquitylated sites are shown in SFigure 10A, 10B and previously described sites in SFigure 10C, 10D, 10E. CURTAIN generated bar charts for CPT1A (K508) and TOMM70 (K573) ubiquitylation site levels confirmed an increase across all replicates of AO treated neuron samples and this is also confirmed by CURTAIN generated violin plots for these sites. TOMM70 encodes a key subunit of the Translocase of Outer Membrane (TOM) and consists of multiple repeating units known as TetraTricopeptide repeat (TPR) domains. The N-terminal TPRs form a loosely structured region called the NTD- pocket, which primarily interacts with molecular chaperones e.g. heat shock proteins (Fig 5E). The C-terminal TPRs form the CTD-pocket, which specifically binds to mitochondrial preproteins for import into the mitochondria (Fig 5E). Analysis of an AlphaFold structural model of mouse TOMM70 reveals that the previously reported Parkin-mediated ubiquitylation sites span multiple TPR domains of the NTD ((K171, K172, K181, K188, K240 and K248) and CTD pockets. Interestingly the newly identified TOMM70 site (K573) lies within a common surface in the CTD-pocket forming a distinct ring-like pattern with the previously reported sites of TOMM70 (K473, K504, K539) (Fig 5F).

## Discussion

We generated two unique tools CURTAIN and CURTAIN-PTM is that it can be used by a non-MS-expert to explore complex MS-based quantitative proteomic and PTM data. These are free to use, open-source, and web-based software requiring no installation. Both tools enable any end user to freely share and explore MS-based proteomic data with a unique weblink. To our knowledge there are no such free to use open-source software packages similar to CURTAIN and CURTAIN-PTM. The programs that are closest to CURTAIN are SimpliFi^™^ (https://protifi.com/pages/simplifi) (33) and Mass Dynamics (https://massdynamics.com) (34) that are commercial web-based tools capable of generating volcano plots and displaying deconvoluted primary data. These commercial packages lack several of the features built into CURTAIN most importantly allowing data to be freely shared and analyzed by any end user.

Moreover, they do not perform PTM analysis and lack the ability to link to curated pathway components of interest such as kinases, phosphatases, ubiquitin components of custom protein sets. They also do not link to protein domain, Alphafold structure or other external sites that CURTAIN and CURTAIN-PTM links to. It should be noted that CURTAIN and CURTAIN-PTM do not store the primary MS raw data and thus should be considered as a companion but not replacement to the PRIDE database (26). For immunological proteomic MS data, the ImmPRes (Immunological Proteome Resource, http://immpres.co.uk/) is widely used (35). This operates as a curated database with interactive visualization of data, but there is no straightforward way to upload your own data to this and share links to enable others to explore their data and deconvolute and analyze your own proteomic data with a sharable weblink. The ImmPRes database does not currently allow the analysis of PTM data.

CURTAIN-PTM is currently compatible with PTM data derived from MaxQuant search that maps the modified sites on peptides along with the site probability information and quantification. The search output for PTM data from other search algorithms such as MS-Fragger, Spectronaut, and DIA-NN are not directly fully compatible with CURTAIN-PTM as this displays the data differently from MaxQuant. For example, the output from Spectronaut displays the modified peptide without information on the residue number of the protein that it is derived from. Unlike MaxQuant, Spectronaut does not output separate PTM files for each modification and only provides a combined file that lists all of the modifications. DIA-NN provides the PTM site probability files and quantification data as different files which would need to be merged and processed prior to uploading into PTM. The output from MS-Fragger search (DDA-LFQ and TMT, does not do DIA) provides site modification of each modified peptide using number of the protein that it is derived from, but unlike MaxQuant does not provide residue number of peptide that is modified which is required by CURTAIN-PTM. To circumvent this, we have developed a script termed MS-Fragger-CURTAIN-PTM CONVERTER (https://doi.org/10.5281/zenodo.8138524), that identifies peptide modification number allowing the MS-Fragger output to be compatible with CURTAIN-PTM. To make CURTAIN-PTM compatible with Spectronaut and DIA-NN, custom scripts would be required, and we plan to implement these in future versions of CURTAIN-PTM.

MS-based proteomic and PTM data is normally generated and shared between collaborators and presented in journals in a static format either in Figures and/or Tables linked to the PRIDE database that hosts the raw MS data (26). These formats, especially for non-MS expert users’, can be challenging to download and explore. This can result in MS-datasets not achieving their potential impact as crucial data that could be discovered and exploited by other researchers, remains buried within these non-interactive datasets. We believe CURTAIN and CURTAIN-PTM will significantly facilitate the exploration of published MS Proteomics and PTM data and enable quality and robustness of the MS data to be more readily assessed. We therefore advocate that a sharable CURTAIN and/or CURTAIN-PTM weblink or equivalent package, be reported in all figure legends to all figures or tables displaying MS data. We have done this for a recent study (36). CURTAIN and CURTAIN-PTM are also extremely useful in sharing MS data between lab members and collaborators. Altogether, these tools align with the FAIR principles to share experimental data (Findable, Accessible, Interoperable and Reusable), we developed two tools that facilitate open science, open source, and ultimately replication of scientific advances (37).

Lastly, we have specifically designed CURTAIN and CURTAIN-PTM for proteomic and phospho-proteomic data, but envisage adapting CURTAIN to analyzing other kinds of OMIC data including metabolomic, lipididomic, or RNA-Sequencing data, therefore expanding the potential impact of these analytical tools to different research fields and human diseases.

In conclusion, CURTAIN and CURTAIN-PTM are free to use open-source useful tools, enabling, sharing and exploration of complex MS-based proteomics data. These are designed to be used by non-MS-experts. We recommend that published studies involving proteomic experiments contain a shareable web-link, allowing reviewers and readers to explore the data interactively.

This will maximize the benefit and impact of published proteomics data.

## Materials and Methods

### Materials

**Table 1.**

| Protein | Company | Catalog number | RRID number | Antibody species | Dilution Used |
| --- | --- | --- | --- | --- | --- |
| *GAPDH* | Santa Cruz | sc-32233 | AB_627679 | Mouse | IB (1:2000) |
| ***LRRK2***<br>***Cterm*** | **NeuroMab** | **N4241A/3**<br>**4** | **AB_2877**<br>**351** | **Mouse** | **IB (1:1000)** |
| ***LRRK2***<br>***pSer935*** | **MRC PPU**<br>**Reagents**<br>**and**<br>**Services** | **UDD2** | **AB_2921**<br>**228** | **Rabbit** | **IB (1:1000)** |
| ***PPM1H* (1-**<br>**514)** | **MRC PPU**<br>**Reagents**<br>**and**<br>**Services** | **DA018** | **AB_2923**<br>**281** | **Sheep** | **IB (1 µg/ml)** |
| ***PPM1H*** | **Abcam** | **ab303536** | **AB_2941**<br>**812** | **Rabbit** | **IB (1 µg/ml)** |
| ***Rab10***<br>***pThr73*** | **Abcam** | **ab230261** | **AB_2811**<br>**274** | **Rabbit** | **IB (1:1000)** |
| ***Rab10 total*** | **nanoTools** | **0680-**<br>**100/Rab1**<br>**0-605B11** | **AB_2921**<br>**226** | **Mouse** | **IB (1:500)** |
| ***PPM1H***<br>**(*CRISPR*)** | **DU64668** | **PPM1H C-term KI BD-TAG-IRES2-**<br>**GFP Sense guide A** |  |  | **pBabeD** |
| ***PPM1H***<br>**(*CRISPR*)** | **DU64674** | **PPM1H C-term KI BD-TAG-IRES2-**<br>**GFP Antisense guide A** |  |  | **pBabeD** |
| ***PPM1H***<br>**(*CRISPR*)** | **DU69480** | **PPM1H C-term KI BD-TAG-IRES2-**<br>**GFP Donor** |  |  | **pMK-RQ** |

### Methods

CURTAIN was created using the Angular Web framework (https://angular.io/). The code is freely available on Github for CURTAIN (accession number https://doi.org/10.5281/zenodo.8138357), CURTAIN-PTM (accession number https://doi.org/10.5281/zenodo.8138360) and associated components, namely backend (accession number https://doi.org/10.5281/zenodo.8138456), UniprotParserjs (accession number https://doi.org/10.5281/zenodo.8138494), common access

Application Protocol Interface (API) allowing others to link to and interact with CURTAIN backend (accession number https://doi.org/10.5281/zenodo.8138473) and MS-Fragger-CURTAIN-PTM CONVERTER (accession number https://doi.org/10.5281/zenodo.8138524) via the MIT open-source license (https://opensource.org/license/mit/). CURTAIN is designed to run on all commonly used web browsers and has been tested on Safari, Chrome, Microsoft Edge and Firefox.

### CURTAIN frontend implementation

The workflow of CURTAIN is described in Figure 1A. The input selection parameters for CURTAIN are selected through an inbuilt dropdown menu (Step 3, Fig 1A) and summarized in (Dataset S1). Experimental metadata can be added in the same format as used by the PRIDE database (26) (Step 3, Fig 1A). Retrieval of up to date Uniprot metadata (Step 4, Figure 1A) uses the Uniprot web API (22). Visualization of the quality controls profile plot and global correlation matrix (Step 5, Fig 1A), Volcano Plot (Step 6, Fig 1A), bar graphs/violin plots (Step 7, Fig 1A), protein domain structure, protein expression profile (Step 8, Fig 1A) employs Plotly (https://plot.ly) (38). In Step 8, Figure 1A, domain structures, disease links, mutagenesis information and pharmaceutical use are determined using Uniprot (https://www.uniprot.org/) (22), predicted structure using AlphaFold (https://alphafold.ebi.ac.uk/) (27, 39), protein interactions deploying the STRING interaction database (https://string-db.org/) (28) as well as the Interactome Atlas (http://www.interactome-atlas.org/) (29) and tissue expression uses ProteomicsDB (https://www.proteomicsdb.org/) (30).

### CURTAIN-PTM frontend implementation

CURTAIN-PTM is designed using the same framework described above. The workflow of CURTAIN-PTM is described in Figure 1B. Steps 1 to 8 in CURTAIN-PTM are the same as CURTAIN. The input selection parameters for CURTAIN-PTM are selected through an inbuilt dropdown menu (Step 3, Fig1B) and summarized in (Dataset S1). The visualization of experimentally determined PTM in each protein (Step 10 Fig 1B) and comparison of experimental and database PTMs (Step 13, Fig 1B) is undertaken using Plotly (https://plot.ly) (38). For predictions of kinases that phosphorylate a phosphorylation site of interest (Step 11, Fig 1B) we deployed the web API from kinase library (https://kinase-library.phosphosite.org/site) (31). For selection of PTM databases (Step 12, Fig 1B) we used PhosphoSitePlus (https://www.phosphosite.org/) (18), Protein Lysine Modification Database (PLMD) using downloaded data (19), CarbonylDB (http://carbonyldb.missouri.edu/CarbonylDB/index.php/) (20), GlyConnect (https://glyconnect.expasy.org/) (21) or Uniprot (https://www.uniprot.org/) (22). For PTM Motif analysis (Step 9, Fig 1B) we used weblogo (https://logojs.wenglab.org/app/) (40). For multiple sequence alignments in CURTAIN-PTM we used biomsalign (https://github.com/ppillot/biomsalign).

### CURTAIN and CURTAIN-PTM backend implementation

We designed CURTAIN and CURTAIN-PTM so that the end user can save their session (Step 9 CURTAIN, Fig 1A, Step 13 CURTAIN-PTM, Fig 1B) and generate a unique web link to retrieve and share the analysis of data and plots, which can also be published (Step 10 CURTAIN, Fig 1A, Step 14 CURTAIN-PTM, Fig1B). If the user has logged into CURTAIN or CURTAIN-PTM employing an ORCID (https://orcid.org/), this associates all data generated by that user and they will be able to view and track all saved sessions associated across different projects within an ownership table (Step 11 CURTAIN, Fig 1A, Step 15 CURTAIN-PTM, Fig 1B). By default when a user has logged in using an ORCID, the data is saved as private. For data to be viewed and shared by others with a weblink, the setting needs to be manually changed by the owner to “Share” (Step 11 CURTAIN, Fig 1A, Step 15 CURTAIN-PTM, Fig 1B). If the user does not login with a user ID then all data can be viewed and shared with anyone through the saved web link.

Within the backend, the user information table contains all the basic non-session related data such as username and password. When the user chooses to save their working session, the frontend will package the current input data, plot settings and user data selections into a single JavaScript Object Notation (JSON) (41) and send the data to the backend. The overall flow of data is summarized in (SFig 1A). The backend was developed using the Python-based Django web framework (https://djangoproject.com) to receive and store user data within a Postgres (https://www.postgresql.org) database with a general schema as represented in SFigure 1B. The schema includes five separate tables namely, user information, session ownership, session metadata, session access token, and user data filter list (SFig 1B).

Upon receiving the uploaded session packages, the data is saved into the session metadata table with a unique random string as weblink id. This table contains information on the date the session has been created, unique id, private or share status, and type of CURTAIN session (i.e., total proteomics or PTM). For users’ logging in using ORCID, the backend automatically creates an account using the ORCID as username and an encrypted randomized password.

### Session data retrieval

When viewed through the web browser, CURTAIN first verifies whether the session is marked as “shared” and permission denied if the session is marked “private”. With successful handshake, CURTAIN then requests the associated session data from the backend which is unpacked and processed with the original settings that were saved by the owner. The data that can be viewed and reanalyzed. The new session can be saved as needed and this will generate a new session link rather than overwrite the previous sessions.

### CURTAIN and CURTAIN-PTM backend host server

CURTAIN and CURTAIN-PTM is currently hosted on a dedicated web server, using the Debian 11 operating system. Each component of the backend including SQL database and Django instance, are operated within their own docker container. The frontends are hosted within GitHub Page. We have also provided instruction for self-hosting the frontends using docker and nginx. All the necessary instructions for setting up CURTAIN can be found at https://github.com/noatgnu/CURTAIN.

Note that for the phosphositePlus PTM database that can be optionally selected to compare experimental data within CURTAIN-PTM (Step 11, Fig 1B) requires license compliance for non-academic users’. The minimum recommended hardware requirements for CURTAIN or CURTAIN-PTM is 4 GB of RAM, and 1 TB of SATA HDD allowing groups to self-host.

### Generation of A549 PPM1H-BromoTag CRISPR/CAS9 knock-in cell line

CRISPR was performed using a paired nickase approach to generate A549 PPM1H-BromoTag knock-in cell line. Complementary oligos for the optimal guide pair A (a sense guide 5‘- GAAATGGCCCAGGGGATTGGG and an anti-sense guide 5’- GAGCTTGTTTCCATGTATTAA) were designed to target the C-terminus of PPM1H locus (ENSG00000111110). The sense guide was cloned into the pBabeD P U6 plasmid, and the anti-sense guide was cloned into the pX335 plasmid. The donor DNA containing IRES2 GFP for cell sorting purpose, was cloned into the pMK-RQ plasmid. CRISPR was done by co-transfecting wild-type A549 cells (80 % confluency, 6-well plates) with 500 ng of donor plasmid and 250 ng of each sense and anti-sense plasmids using Lipofectamine LTX according to the manufacturer’s instructions (Life Technologies). The transfected cells were grown for 24 h before selection with fresh DMEM media supplemented with 3 μg/ml puromycin for another 48 h. Single cells with positive GFP signal were sorted, then grown in individual wells of a 96-well plate for approximately 2 weeks. After reaching around 80 % confluency, individual clones were transferred into 6-well plates and homozygous A549 PPM1H-BromoTag clones determined by immunoblotting and genomic sequencing.

### Sample Processing

A549 PPM1H-BromoTag cells were cultured on 15 cm dishes (2 dishes per replicate). Cells were treated with 300 nM AGB1 or cis-AGB for 4 h or 24 h, from a 1000x stock made up in DMSO. Cells were lysed in 600 μl of Lysis Buffer [50 mM Tris–HCl pH 7.5, 150 mM NaCl, 10% glycerol, 10 mM 2-glycerophosphate, 10mM sodium pyrophosphate, 1 mM sodium orthovanadate, 1 μg/ml microcystin-LR, complete EDTA-free protease inhibitor cocktail (Roche) and 1% (v/v) Triton X-100] as described previously (23), and lysates from the 2 dishes if cells pooled in 2 ml low-binding Eppendorf tubes. A small aliquot was taken for immunoblotting and the remainder of the sample was supplemented with 2% (w/v) SDS and subjected to sonication using a probe sonicator (BRANSON) employing 3 pulses. Samples were clarified at 17,000 x *g* at room temperature for 20 min and protein concentration measured using the BCA (bicinchoninic) protein assay (42).

### Sample preparation for total and Phosphoproteomic analysis

3 mg of SDS lysate was allotted for each sample and subjected to S-Trap assisted on-column digestion. The lysate from each sample was reduced by making up to a final 10 mM TCEP (tris(2-carboxyethyl)phosphine) from a 0.1M stock in 300 mM TEAB buffer (Triethylammonium bicarbonate, pH 8.0) and incubated on a Thermomixer at 60°C for 30 min with an agitation set at 1200 rpm. Samples were then brought to room temperature and alkylated by adding 40 mM Iodoacetamide (IAA) and further incubated on a Thermomixer at room temperature for 30 min with an agitation set at 1200 rpm and then quenched by adding additional further 5 mM TCEP and incubated at room temperature for 10 min. From a stock of 20% (w/v) SDS prepared in milliQ water, SDS concentration of each sample was increased to 5% (w/v) and acidified by bringing the final concentration to 1.2% (v/v) phosphoric acid from a 12% stock prepared in milliQ water. Immediately, 7 times the volume of lysate, S-Trap buffer (90% methanol (vol/vol) in 100 mM TEAB buffer) was added, and the resulting solution transferred to a S-Trap midi column. The columns were centrifuged at room temperature at 2000 rpm for 1 min. Columns were washed by adding 4 ml of S-Trap buffer and centrifuged at 2000 rpm for 1 min and this step was repeated four times. Flowthrough was discarded at each step and after the final wash, the columns were centrifuged at 3000 rpm for 1 min and the columns were transferred to a new 15 ml collection tubes. For each S-Trap column, 30 µg (1 to 100 ratio of protease to cell extract amount) of a mixture of Trypsin+Lys-C (Thermo A41007) dissolved in 350 µl of 50 mM TEAB and was added. In addition, 300 µg of Tosyl phenylalanyl chloromethyl ketone (TPCK) treated trypsin (Sigma,4352157) was dissolved in 350 µl of 50 mM TEAB buffer and added to the S-TRAP column. Columns were immediately centrifuged at 100 rpm for 1 min to remove air-bubbles and the flowthrough was re applied to the column which were then placed on a Thermo mixer with a 15 ml rack and incubated at 47 °C for 90 min without agitation. The temperature was reduced to room temperature and digestion was continued the on-column digestion overnight (∼ 16 h). 500 µl of 50 mM TEAB was then added and column centrifuged 100 rpm for 1 min. Next, 500 µl of 0.1% (v/v) formic acid in Milli-Q water was added and centrifuged 100 rpm for 1 min. 500 µl of 80 % (v/v) Acetonitrile in 0.1% formic acid (v/v) was added and centrifuged at 100 rpm for 1 min and this step two further times. Eluates were then vortexed and centrifuged at 3,000 rpm for 10 min to precipitate any debris or undigested protein amounts. Eluates were then transferred to 2 ml low binding Eppendorf tubes, placed on dry ice for 10 min and vacuum dried using a speedvac concentrator and if required stored in -80 freezer.

The dried samples were resuspended in 500 µl 1% (v/v) TFA (trifluoroacetic acid) and incubated for 30 min on a thermomixer at 1800 rpm at room temperature, sonicated on a water bath sonicator for 10 min and centrifuged at 17,000 x *g* for 10 min at room temperature. The supernatant was transferred to new 2 ml low binding Eppendorf tubes and pH was checked by pipetting 0.5 µl onto the pH strip to ensure samples were pH 2.0. Sep-Pak cartridges (50 mg tC18) were placed into 15 ml falcon tubes. The cartridges were activated by adding 1 ml of 100% acetonitrile and centrifuged at 100 x *g* for 1 min and this step repeated a further 3 times. Cartridges were then equilibrated by undertaking 4 x 1 ml washes with 0.1% (v/v) TFA. Dissolved peptide samples were loaded onto the equilibrated cartridges and allowed to pass through by gravity (takes 15-30 min) and the flowthrough was reapplied. The cartridge was washed four times in 1 ml 0.15% (v/v) formic acid and flowthrough disregarded. Peptides were eluted using 3 washes of 300 µl of 50% acetonitrile in 0.15% (v/v) formic acid. 1 µl aliquots were taken for a digestion check and 10% (90 µl) of each sample was taken for total proteomic analysis. The remaining samples were snap-frozen on dry ice, vacuum dried and stored at -80 °C. A detailed protocol containing this step for combined total and phosphoproteomic analysis is available on Protocols.io (**<u>dx.doi.org/10.17504/protocols.io.261ged49yv47/v1</u>**).

### TiO2-based Phosphopeptide enrichment

Phospho-peptide enrichment was carried out using the High-Select TiO_2_ Phospho-peptide Enrichment kit (Thermo Fisher, A32993), as per manufacturer’s instructions. Lyophilized samples were resuspended in 150 µl of “Binding/Equilibration buffer” (provided with the kit) and incubated at room temperature on a Thermomixer at 1800 rpm for 30 min. Samples were sonicated in a water bath sonicator for 10 min and centrifuged at 17,000 x *g* for a further 10 min. The supernatant was transferred to a protein low-binding 1.5 ml Eppendorf tube and pH was checked by pipetting 0.5 µl onto the pH strip to ensure samples were <pH 3.0. TiO_2_ spin tip (provided in kit) was placed into a 2 ml low binding Eppendorf tube. Tips were washed using 20 µl wash buffer (provided in kit) and centrifuged at 3000 x *g* for 2 min, and equilibrated in 20 µl Binding/Equilibration buffer and centrifuged for a further 2 min. The equilibrated tip was transferred to a new 2 ml protein low-binding Eppendorf tube. The resuspended samples were loaded onto the tip and centrifuged at 1000 x *g* for 5 min. The flowthrough was re-applied, and centrifugation repeated. The tips were transferred to a new 2 ml protein low-binding tube, washed with 20 µl Binding/Equilibration column, followed by 20 µl wash buffer, centrifuging at 3000 x *g* for 2 min each time. Both wash steps were repeated one more time and washed with 20 µl of LC-MS grade water. The tips were placed in a new 1.5 ml protein low-binding Eppendorf tube and the phospho-peptides eluted using 50 µl Elution buffer (provided with kit) and centrifugation at 1000 x *g* for 5 min. The Elimination step was repeated one more time. 2 µl was taken for a phospho-enrichment check. The remaining eluate was snap-frozen on dry ice and vacuum dried. Samples were then subjected to C18 clean-up using 50 mg 1cc tC18 Sep-Pak cartridges, as described above. A detailed protocol containing this step for combined total and phosphoproteomic analysis is available on Protocols.io (**<u>dx.doi.org/10.17504/protocols.io.261ged49yv47/v1</u>**).

### TMT labeling

The Phospho-enriched as well as the total proteomic samples were resuspended in 30 µl fresh 50 mM TEAB and incubated at room temperature on a thermomixer at 1800 rpm for 1 hr. Samples were centrifuged at 17,000 x *g* for 3 min and supernatant transferred to 0.5 ml protein low binding Eppendorf tube. 0.5 mg of each TMT reagent (Thermo Fisher, A44520) was resuspended in 100 µl 30% anhydrous acetonitrile to a final concentration of 5 µg/µl. TMT labels were added to the samples (10 µl for phospho-proteomics, and 20 µl for total proteomics), incubated at room temperature on a thermomixer at 1200 rpm for 2 h and 50 mM TEAB added to adjust the total volume to 100 µl. The mixtures were incubated on a thermomixer at 1200 rpm for a further 10 min at room temperature. A total of 2 µl aliquots of each sample were removed to verify the TMT labeling efficiency. Furthermore, 2 µl of each sample was removed and pooled to prepare a mini-pool to verify the absolute peptide abundance in each sample. Once label checks were complete, samples were quenched by adding 3 µl of 5% hydroxyl amine and incubating at room temperature on a thermomixer at 1250 rpm for 20 min. Samples were then pooled appropriately to achieve equal abundance of each TMT labeled peptides. We have generated a simple web-based tool to determine the specific volumes of each sample that need to be mixed together to equalize the peptide levels (https://samplepooler.proteo.info). The final mixture was centrifuged at 17,000 rpm for 1 min and, snap-frozen on dry ice and vacuum dried. A detailed protocol containing this step for combined total and phosphoproteomic analysis is available on Protocols.io (**<u>dx.doi.org/10.17504/protocols.io.261ged49yv47/v1</u>**).

### High-pH reversed-phase liquid chromatography fractionation

Total proteome and phosphoproteome samples were next fractionated using high-pH reversed-phase liquid chromatography fractionation as described previously (43). Dried samples were resuspended in 110 µl Solvent A [10mM ammonium formate in LC-MS grade water, pH adjusted to 10.0 using MS grade ammonium hydroxide] and incubated at room temperature on a thermomixer at 1800 rpm for 20 min. 100 µl of the phospho-proteomic and total proteome sample was transferred to LC vials for high pH fractionation using xbridge 25 cm C18 column (Waters, 186003010) on a dionex 3000 LC system. The column was equilibrated at 3% solvent B (10 mM ammonium formate in 80% acetonitrile pH 10) at 0.275 ml/min flow rate. The sample was loaded and the column washed with 3% solvent B for 20 min to remove excess unlabelled TMT reagent. A linear gradient from 3 to 40% solvent over 90 min was applied and 96 fractions were collected in a 96 deep well plate. Fractions were concatenated into 48 protein low binding 1.5 ml Eppendorf tubes (<u>dx.doi.org/10.17504/protocols.io.bs3tngnn</u>), snap-frozen, vacuum dried and stored at -80 °C.

Dried fractions were resuspended in 50 µl LC buffer [3% acetonitrile in 0.5% Formic acid] and incubated on a thermomixer at 1800 rpm for 1 h at room temperature. Samples were vortexed and centrifuged at 17,000 x *g* for 5 min. 10 µl of each sample was transferred to LC vials and subjected to mass spectrometry. A detailed protocol containing this step for combined total and phosphoproteomic analysis is available on Protocols.io (**<u>dx.doi.org/10.17504/protocols.io.261ged49yv47/v1</u>**).

### MS data acquisition and database searches

Phosphopeptides were analyzed on a Orbitrap Lumos Tribrid mass spectrometer in-line with Dionex RSLC 3000 nano-liquid chromatography system. Peptides were loaded on a 2cm trap column at 5 µl/min flow rate and resolved on a 50 cm analytical column at 350 nl/min flow rate. Data was acquired using Data dependent acquisition in MS2 mode. Full MS was acquired in the mass range of 350 - 1500 m/z and measured using Orbitrap mass analyzer at a resolution of 120,000 at m/z 200. MS2 scans were acquired in a top 15 data dependent mode and fragmented using a normalized higher energy collisional dissociation (HCD) 37.5% and measured using Orbitrap mass analyzer at a resolution of 45,000 at m/z 200. Quadrupole mass filter was set to 0.7 Da for MS2 scans. AGC targets and Ion injection times for both MS1 and MS2 were set at 3E6 and 2E5 for 30 ms and 120 ms respectively. Total proteome was analyzed using the same MS instrument and columns except the data was acquired using SPS-MS3 mode. Full MS1 was acquired using an Orbitrap mass analyzer at a resolution of 120,000 at m/z 200. MS scans were acquired at a top speed for 2 sec and fragmented using 32% HCD and measured using ion trap mass analyzer. Synchronous precursor selection (SPS) for MS3 was performed using 10 MS2 fragment ions and further fragmented using normalized 65% HCD energy and measured using Orbitrap mass analyzer at a resolution of 60,000 at m/z 200. A detailed protocol containing this step for combined total and phosphoproteomic analysis is available on Protocols.io (**<u>dx.doi.org/10.17504/protocols.io.261ged49yv47/v1</u>**).

Total proteome raw data was converted to mzML using MS-covert tool (44) and imported into FragPipe version 18.0. In-built TMT-Pro16 plex workflow was selected as default settings and searched data using MSFragger version 3.5 against the Human Uniprot database (July, 2022: UP000005640). Oxidation of Met, Phosphorylation of STY, Deamidation of NQ were selected as variable modifications and Carbamidomethylation of Cys as fixed modifications. Trypsin as protease with two missed cleavages were allowed. Phosphoproteomics data was searched using MaxQuant version 2.0.3.0 and searched against the Human Uniprot database (May 2021). TMT-Pro 16plex workflow was selected with 0.75 Precursor ion filter (PIF) was sent as a filter. Trypsin was selected as protease with a maximum of two missed cleavages were allowed. Oxidation (M);Acetyl (Protein N-term);Deamidation (NQ) and Phosphorylation of (STY) were set as variable modifications and Carbamidomethylation of Cys as fixed modification. 1% False discovery rate for PSM, sites and protein level was applied. The search output files from total proteome and Phosphoproteomics data was further processed using Perseus version (1.6.15.0) for statistical analysis and further data was visualized using CURTAIN and CURTAIN-PTM.

## Acknowledgements

We thank Alessi Team Aligning Science Across Parkinson’s (ASAP) members and Rubén Fernández-Santiago for important feedback and discussion. We thank Hina Ojha for data analysis and preparation of Figure 5E & F, Adam Bond and Alessio Ciulli for providing us with the initial batches of AGB1 and cis-AGB1 and the excellent technical support of the MRC Protein Phosphorylation and Ubiquitylation Unit (PPU) Genotyping team (coordinated by Gail Gilmour), the MRC-PPU DNA sequencing service (coordinated by Gary Hunter), the MRC-PPU tissue culture team (coordinated by Dr Edwin Allen), the MRC-PPU mass spectrometry facility team (coordinated by Dr Renata Soares) and the MRC-PPU Reagents and Services team (coordinated by Dr James Hastie).

## Author contributions

TKP Conceptualization Methodology Investigation Writing—original draft Writing—review & editing

KB Methodology Investigation TLCHBP Methodology Investigation MM Conceptualization review & editing

DRA Conceptualization Methodology Writing—original draft Writing—review & editing

RSN Conceptualization Methodology Investigation Writing—original draft Writing—review & editing

## Data and materials availability

All the primary data that is presented in this study has been deposited in publicly accessible repositories. The CURTAIN and CURTAIN-PTM source code as well as the primary immunoblotting data and statistical analysis have been deposited in Zenodo (https://doi.org/10.5281/zenodo.8182684, https://doi.org/10.5281/zenodo.8182685, https://doi.org/10.5281/zenodo.8138456, https://doi.org/10.5281/zenodo.8183345, https://doi.org/10.5281/zenodo.8138473, https://doi.org/10.5281/zenodo.8138524). Proteomic data have been deposited in the ProteomeXchange PRIDE repository (Identifier:PXD043806). All plasmids and antibodies generated at the MRC Protein Phosphorylation and Ubiquitylation Unit at the University of Dundee can be requested through our website https://mrcppureagents.dundee.ac.uk/. For the purpose of open access, the authors have applied a CC BY public copyright license to all Author Accepted Manuscripts arising from this submission.

## Figure legends

**SFigure 1.**
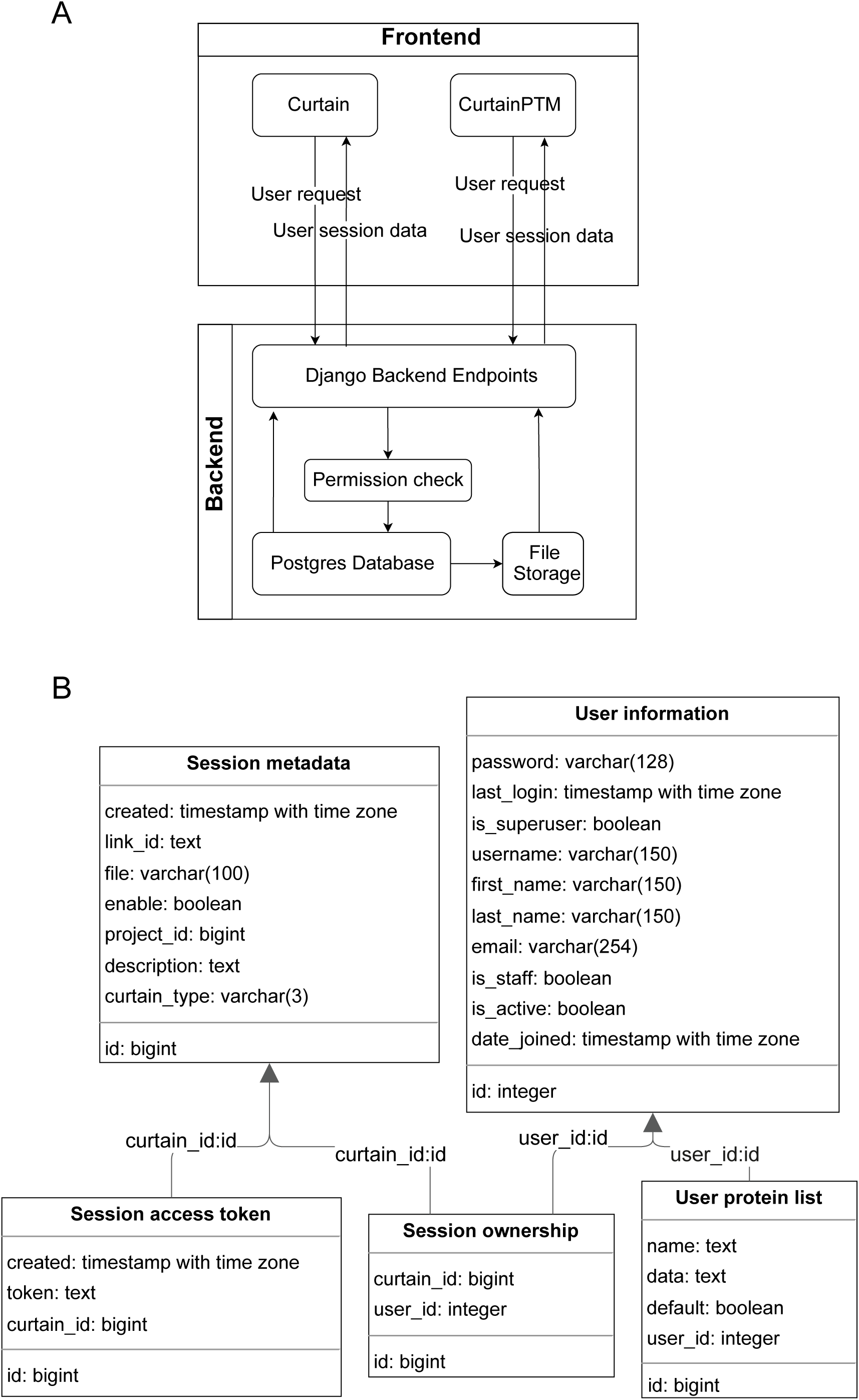
Architecture of CURTAIN and CURTAIN-PTM. (A) CURTAIN and CURTAIN-PTM utilize the same backend. When a user request is accepted by the backend, a permission check is carried out to screen whether the request is sharable or private and can gain access to the database. (B) Summarizes the data structure within the database.

**SFigure 2:**
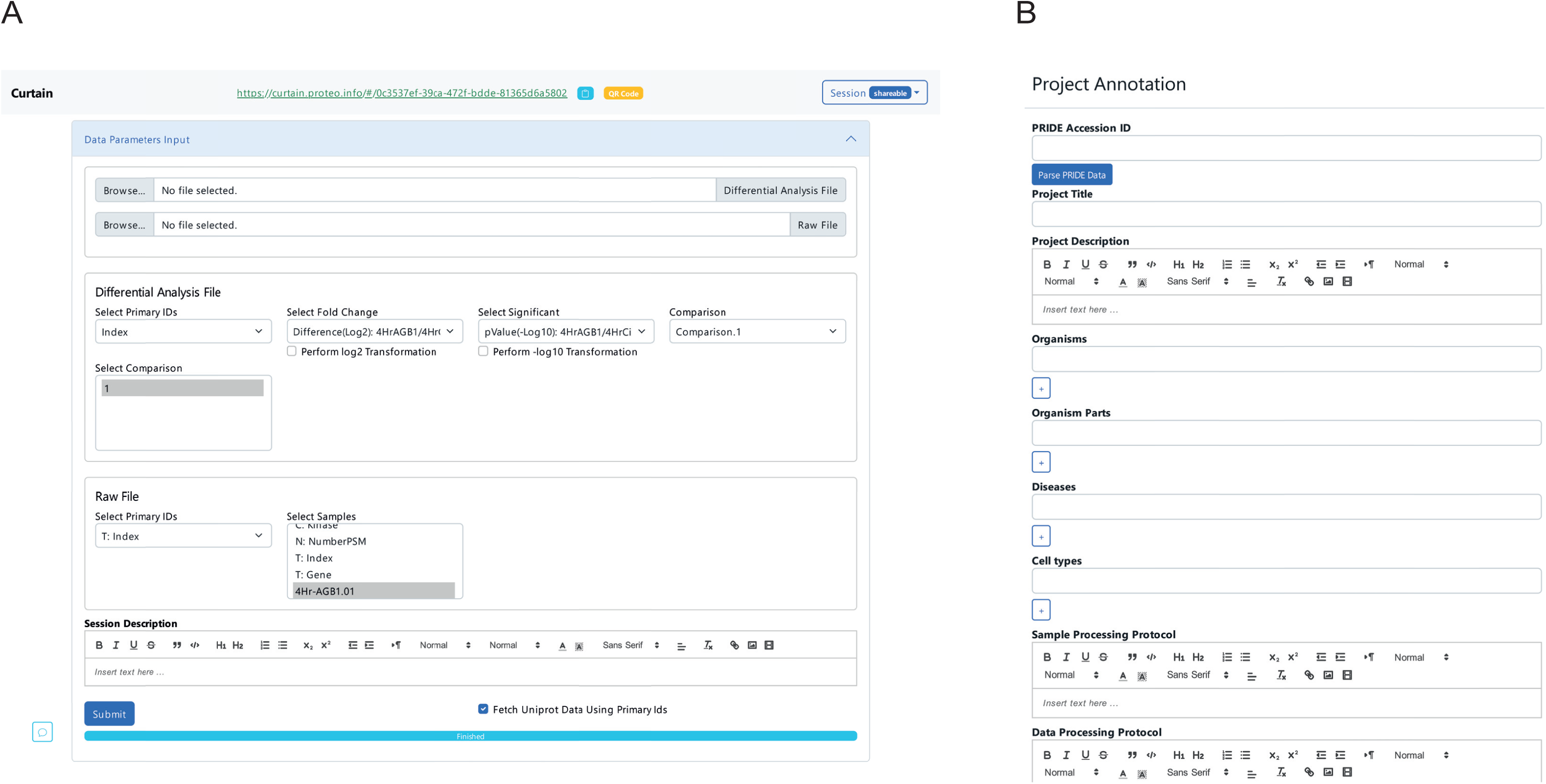
Overview of CURTAIN user interface. (A) Screenshot of the base parameter input interface from a loaded CURTAIN session link. Within the base input, the user can import the two required differential analysis (Differential analysis file from the interface) and processed primary data files (Raw file from the interface). Further detail on selection parameters are described in Dataset S1. From the dropdown under “Session” button, the user can choose to include metadata for the project in Project Annotation (B). The input forms are designed to be similar to that of the data submission process for the PRIDE database.

**SFigure 3:**
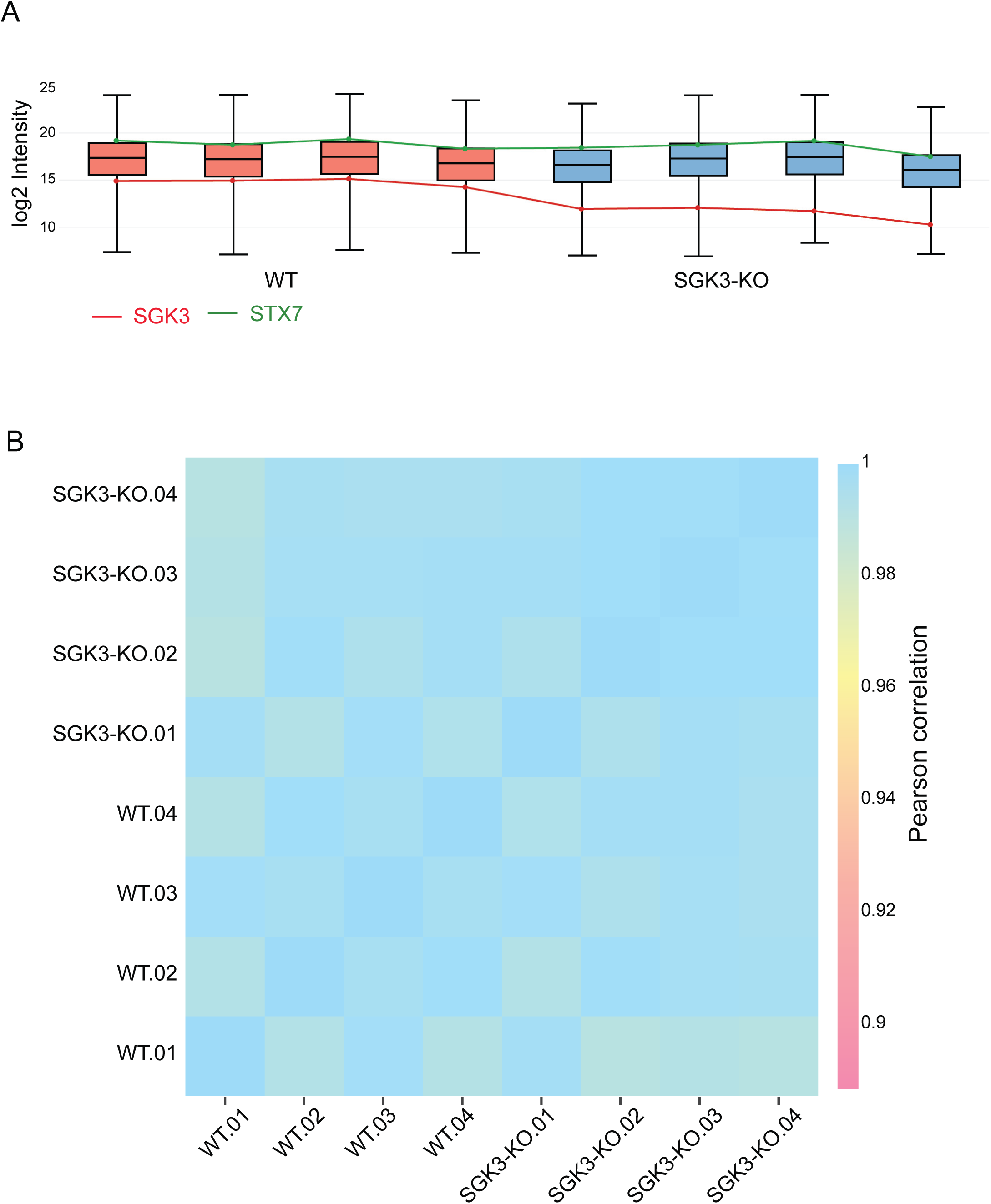
Overview of quality control visualization capabilities of CURTAIN using SGK3 experiment data described in Figure 3. (A) Profile plot that is composed of box plots built from log2 of imported primary data depicting the data distribution and quality of the proteomics data. Within (A), we annotated the primary data from SGK3 and STX7 across all samples. (B) is a correlation matrix calculated within the browser from the same imported primary data. The color scale described the Pearson correlation value within the matrix. The Curtain link for this data is https://curtain.proteo.info/#/6ac93165-d351-4634-9683-ed342a6feaa8

**SFigure 4:**
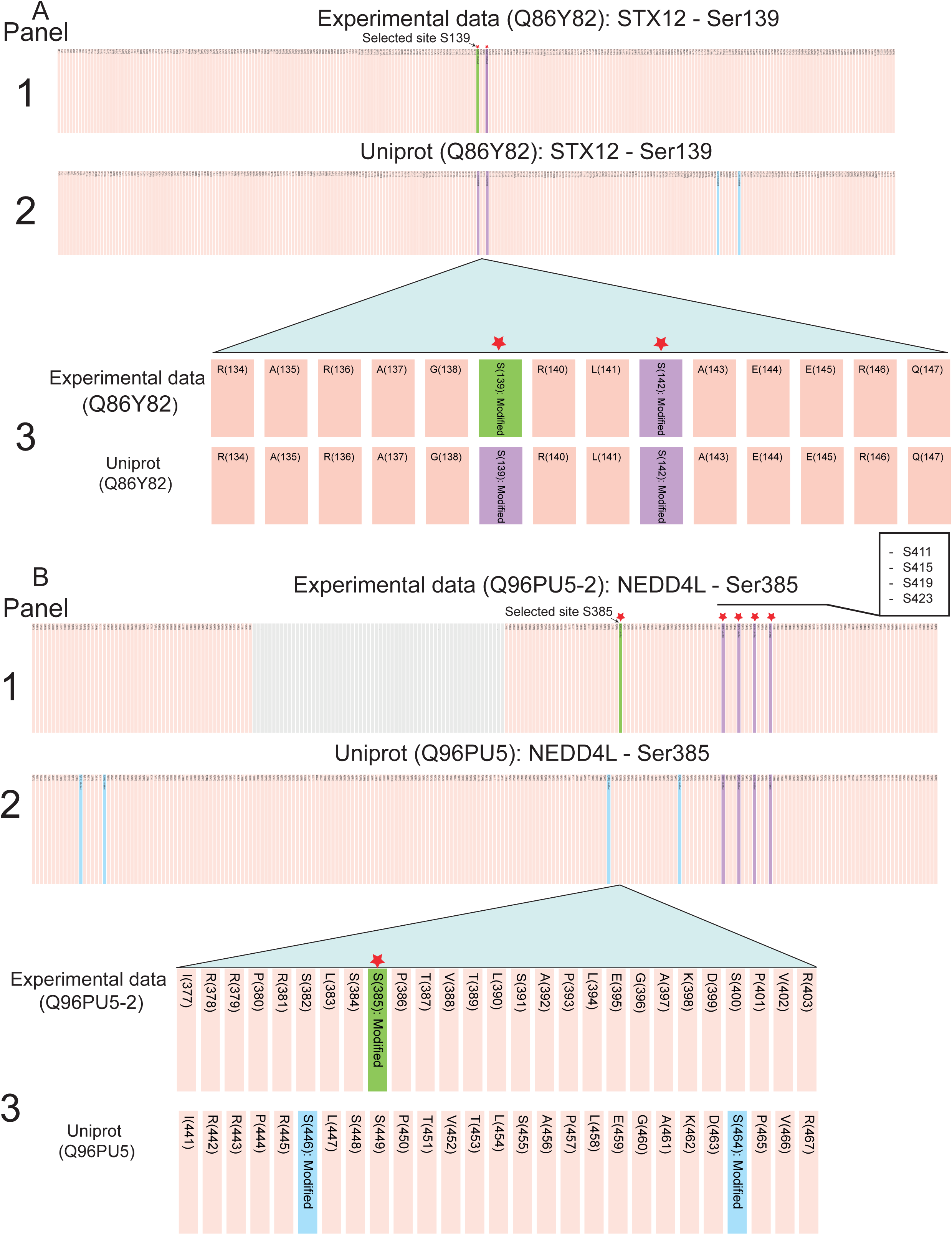
Visualization of PTM data on linear protein sequence. From the analysis undertaken in Fig 4A, two hits namely STX12 and NEDD4L were selected for visualization of PTM data on a linear protein sequence. Note that CURTAIN-PTM numbering of residues is based on “experimental data” provided by the MS search algorithm. which may differ from the canonical sequence displayed in selected databases or discussed in the literature. CURTAIN- PTM analysis of the phosphoproteomic data analyzed in Fig 4A, revealed that STX12 was phosphorylated at 2 sites (Ser139 and Ser142) whilst NEDD4L was phosphorylated at 5 sites (Ser385, Ser411, Ser415, Ser419 and Ser423). The CURTAIN-PTM outputs from these analyses are shown in Panel-1. Highlighted sites selected by the user are in green. Other identified non-selected sites are displayed in purple. CURTAIN-PTM automatically marks STX12-S139 (A) and NEDD4L sites (Ser385, Ser411, Ser415, Ser419 and Ser423) (B) with an asterisk as the levels of sites change significantly between experimental conditions (±IGF1). The other sites not marked with an asterisk do not change with experimental conditions and are not highlighted. Curtain permits experimental PTMs to be compared with publicly available databases such as Uniprot and the output of this comparison is displayed in Panel-2. Other phosphorylation sites that are listed in the database but not detected in the phospho-proteomic data are shown in blue. To better visualize sites of interest, CURTAIN-PTM permits to zoom-in within a selected area Panel-3. The upper panel 3 lists the sequence from the MS search algorithm whilst the lower panel lists the sequence based on the selected database. Note for STX12 (A) the numbering of residues is identical, as the same splice variant was analyzed in MaxQuant and Uniprot, however for NEDD4L (B) the residue numbers differ as Uniprot selected a different isoform to MaxQuant.

**SFigure5:**
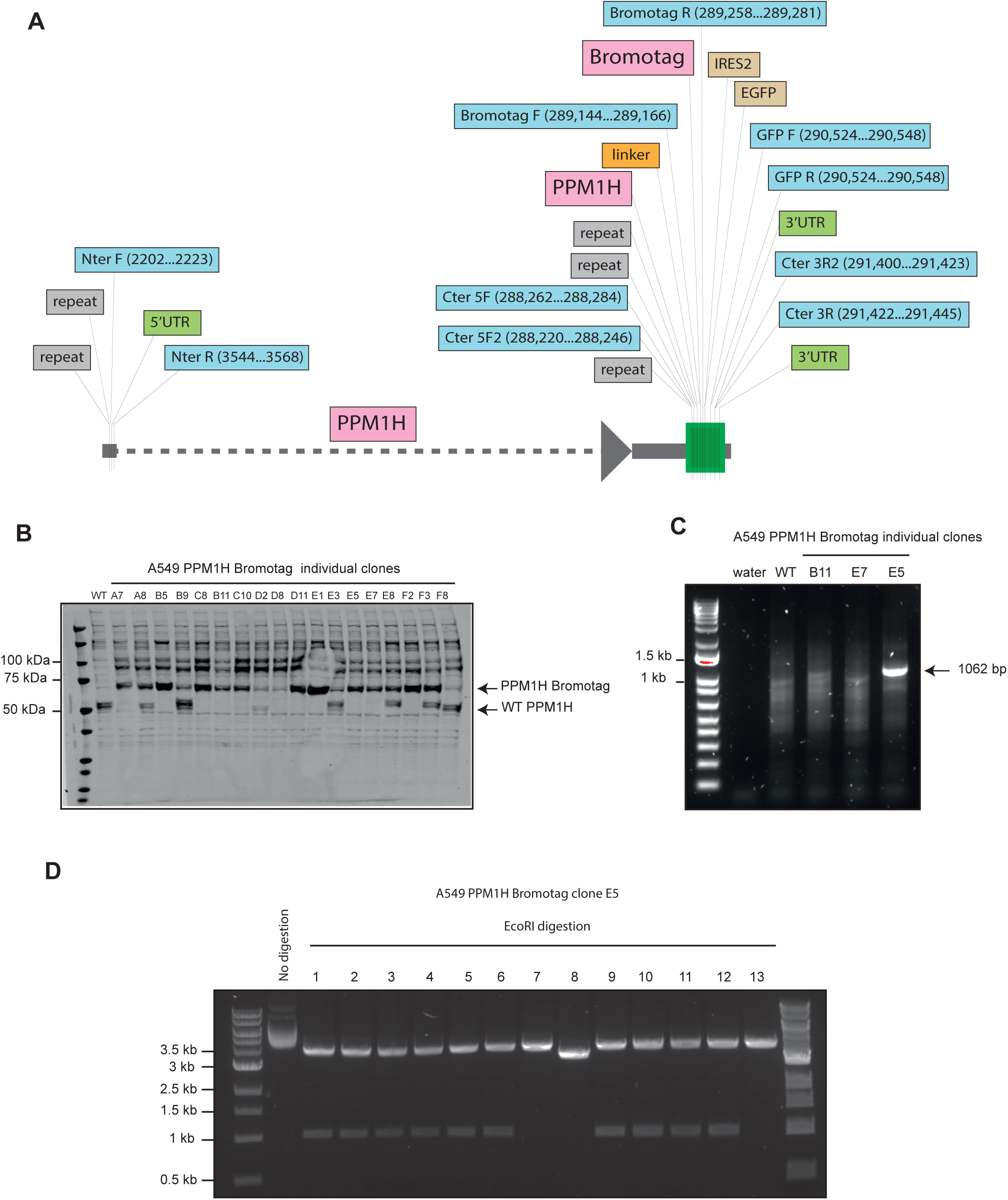
Verification of PPM1H-BromoTag knock-in cells. (A) The map of PPM1H locus containing C-terminal BromoTag and IRES2 GFP. (B) Immunoblotting was performed to screen individual clones of PPM1H-BromoTag using anti-PPM1H antibody at concentration 1 μg/ml (sheep polyclonal antibody, MRC PPU Reagents and Services, DA018). Wild-type (WT) A549 cells were included as a control. (C) PCR was performed to confirm the presence of PPM1H- BromoTag in potential homozygous clones observed from immunoblotting. A pair of primers (Cter 5F2: 5’- CTTGCTGAACTTACATTGGTCAAGAGG and BROMOTAG R: 5’- ACTTGATTGTGCTCATGTCCATGG) were used to amplify an amplicon with expected size at 1062 bp. Water and wild-type (WT) A549 were included as negative controls. (D) The clone E5 was selected to clone into the StrataClone PCR cloning vector pSC-B-amp/kan according to the manufacturer’s instructions (Stratagene). Thirteen random positive colonies were digested with EcoRI restriction enzymes. All thirteen plasmids were sent for sequencing with M13 forward primer (5’- GTAAAACGACGGCCAGTG) and M13 reverse primer (5’- GGAAACAGCTATGACCATG). The plasmids 1, 2, 3, 4, 5, 6, 9, 10, 11, 12 showed sequences matching with the template of PPM1H-BromoTag. In contrast, the plasmids 7, 8, 13 did not show any match with the template, indicating that the fragments cloned into these plasmids are just non-specific PCR products. These results confirm that PPM1H-BromoTag clone E5 is a homozygous clone which was used for all experiments in this study.

**SFigure 6:**
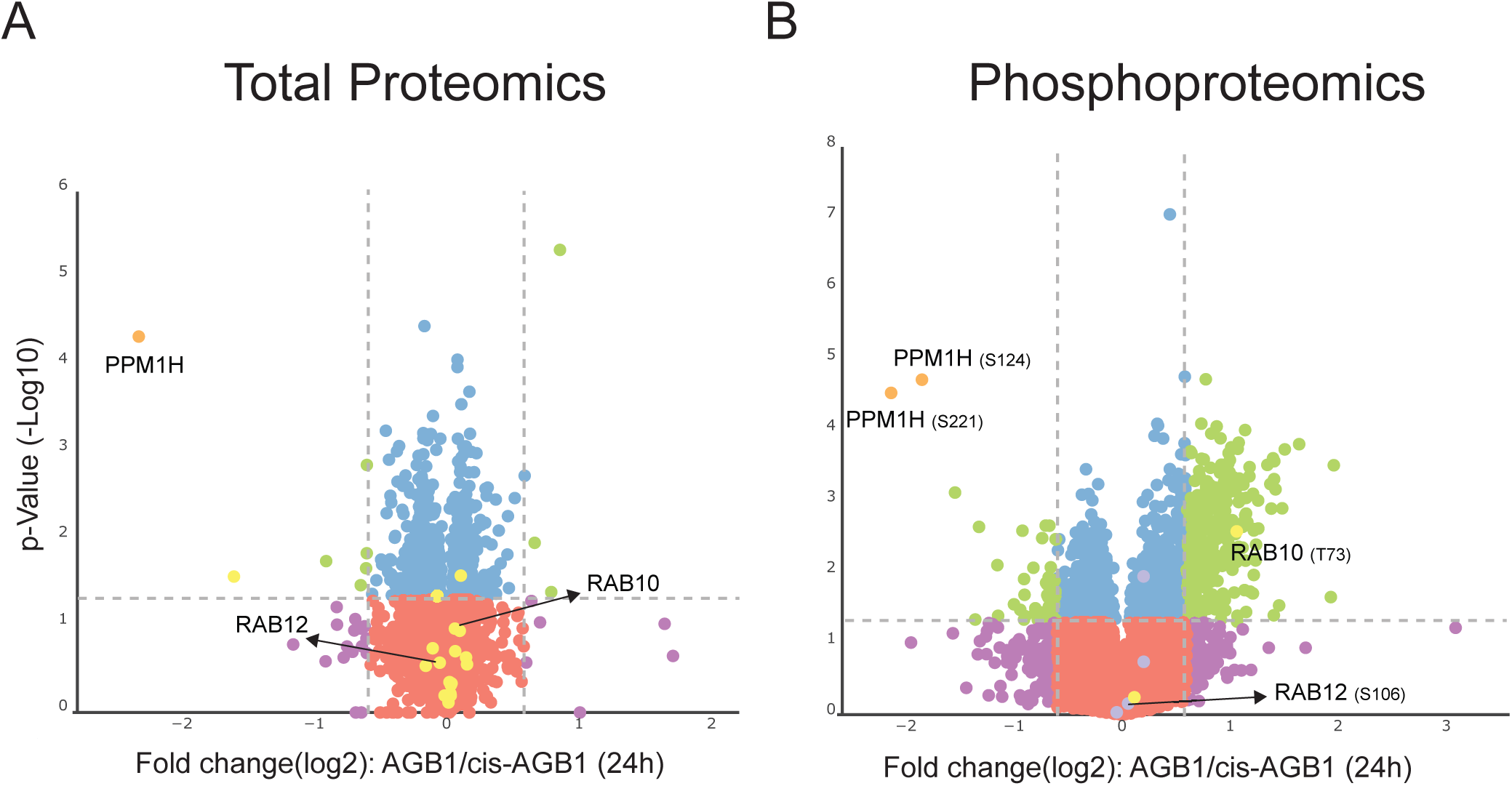
Quality control visualization of PPM1H-BromTAG experimental data shown in Figure 7. (A) Profile plot that is composed of box plots built from log2 of imported primary data depicting the data distribution and quality of the proteomics data. Within (A) we annotated the primary data from total PPM1H and RAB10. (B) is a correlation matrix calculated from the same imported data. The matrix heat map showed that there are minor differences in correlation between 4h and 24h data. The Curtain link for this data is https://curtain.proteo.info/#/273b6d5e-2f21-43a6-a33c-e6ac53e801bd

**SFigure 7:**
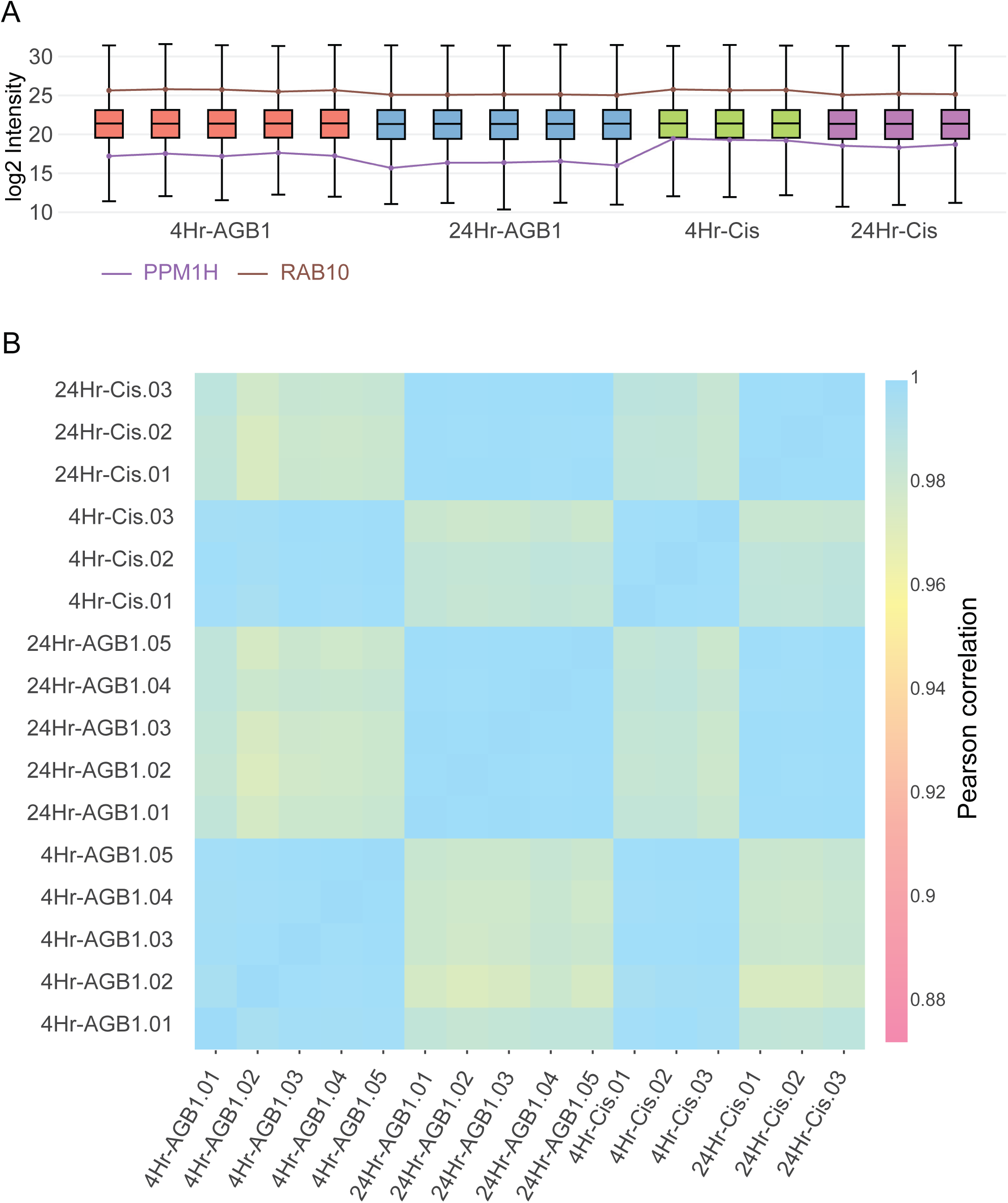
Visualization of PPM1H total and phosphoproteomics data using CURTAIN and CURTAIN-PTM. The lysates produced from the experiments described in Fig 6B in which cells were treated for 24h with cis-AGB1 or AGB1 were subjected to total and phospho-proteomic analysis with data being analyzed with MS-Fragger and MaxQuant. CURTAIN generated volcano plot are presented for the total proteomic (A, https://curtain.proteo.info/#/244cc639-3f22-40e3-b528-564b4989d8f6) and Phospho-proteomic (B, https://curtainptm.proteo.info/#/5510ee39-5695-4995-a085-13adcc47e5cc) and proteins of interest highlighted in black. The same color coding is used as described in Fig 3A. C-E) The primary total and Phosphosite intensities of indicated hits were shown as bar graphs and violin plots.

**SFigure 8:**
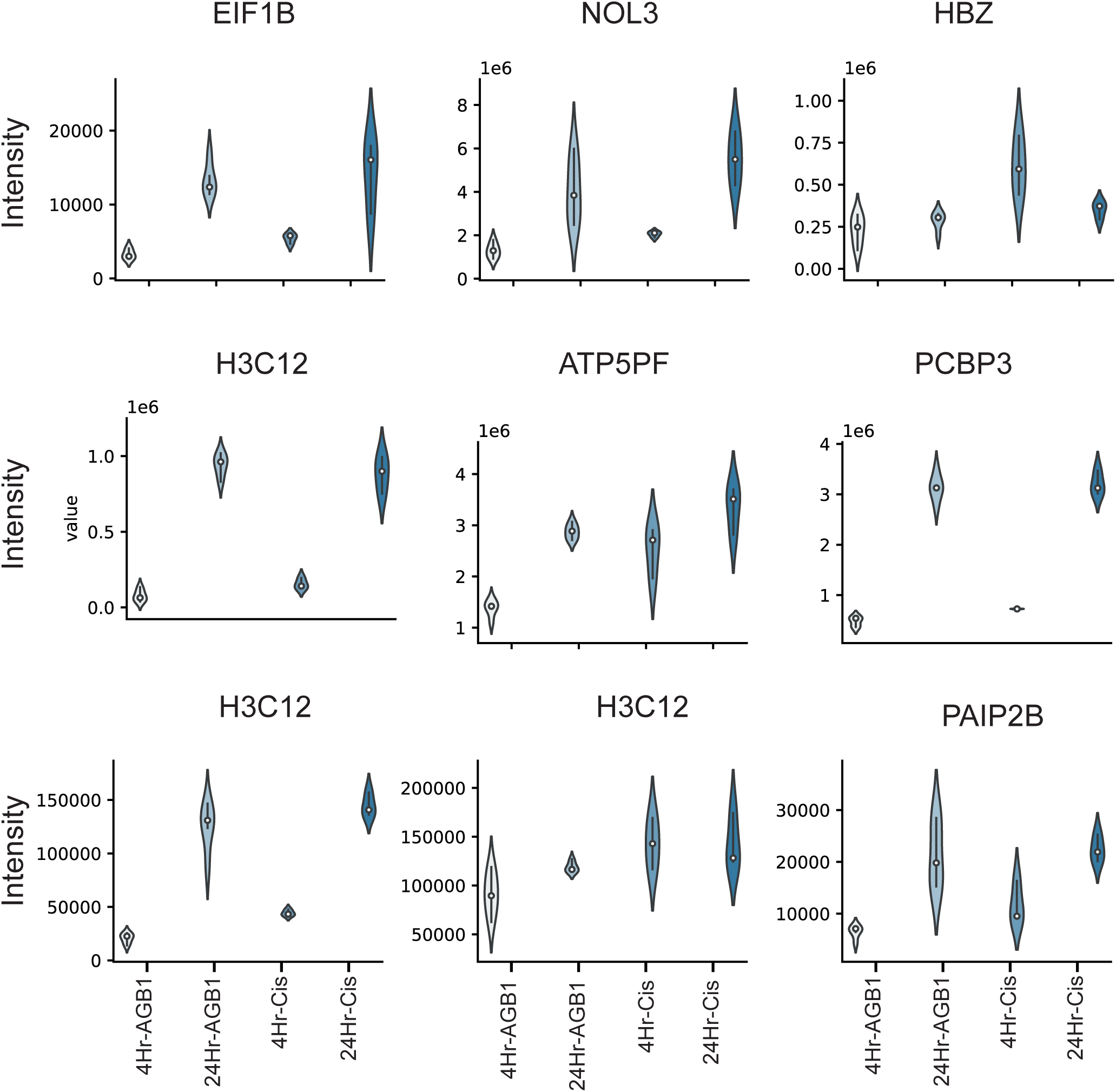
Proteins whose levels are impacted following PPM1H-BromoTag depletion. A grid of violin plots of significantly changed proteins with log2 fold change greater than 1.0 derived from the experiments described in Figure 7A and SFigure 6A. The plots are composed from total proteomics data from both 4h and 24h samples.

**SFigure 9:**
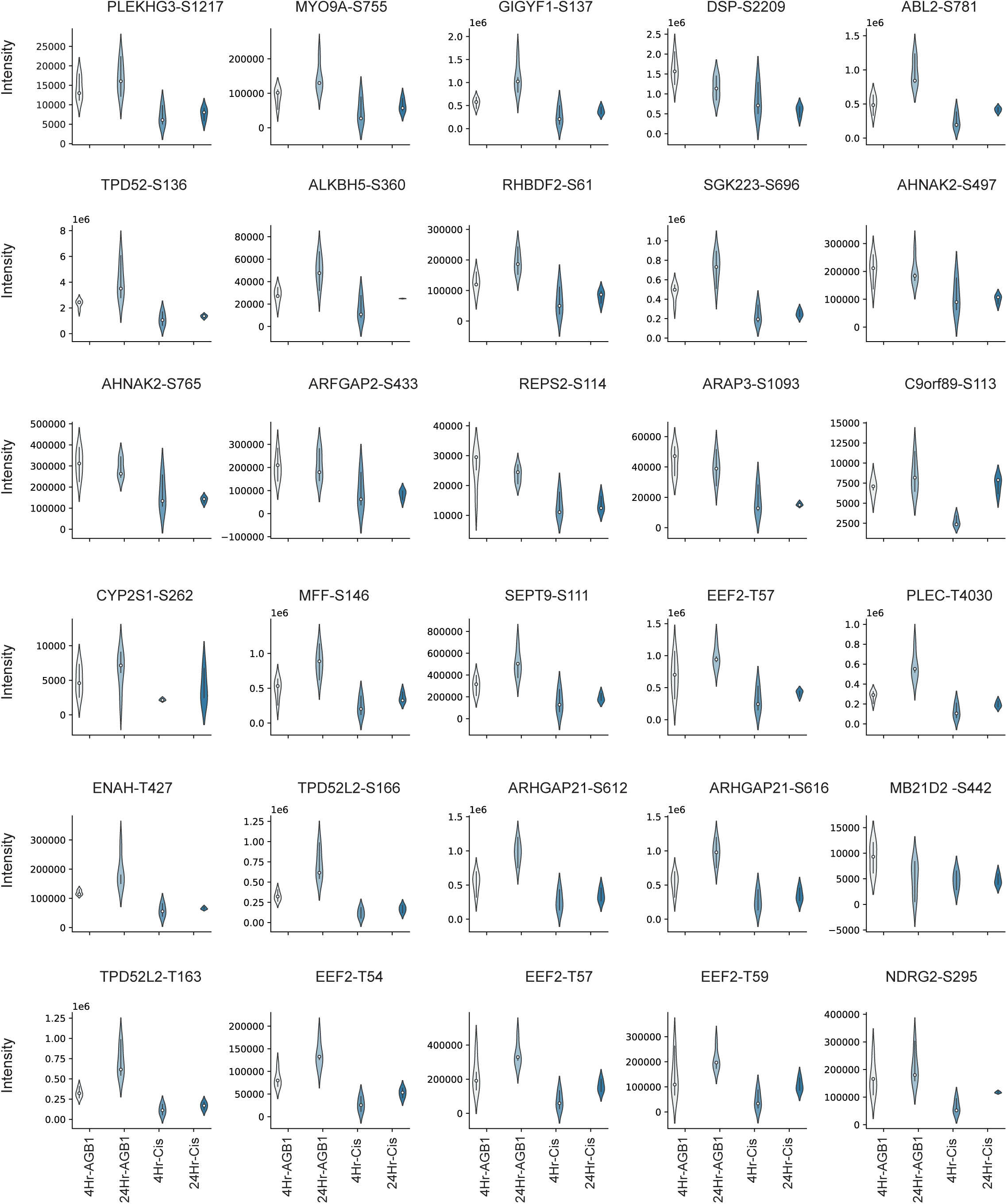
Phosphosites whose levels are impacted following depletion PPM1H- BromoTag. A grid of violin plots of significantly changed proteins with log2 fold change greater than 1.0 derived from the experiments described in Figure 7B and SFigure 6B. The plots are composed from phospho-proteomics data from both 4h and 24h samples.

**SFigure 10:**
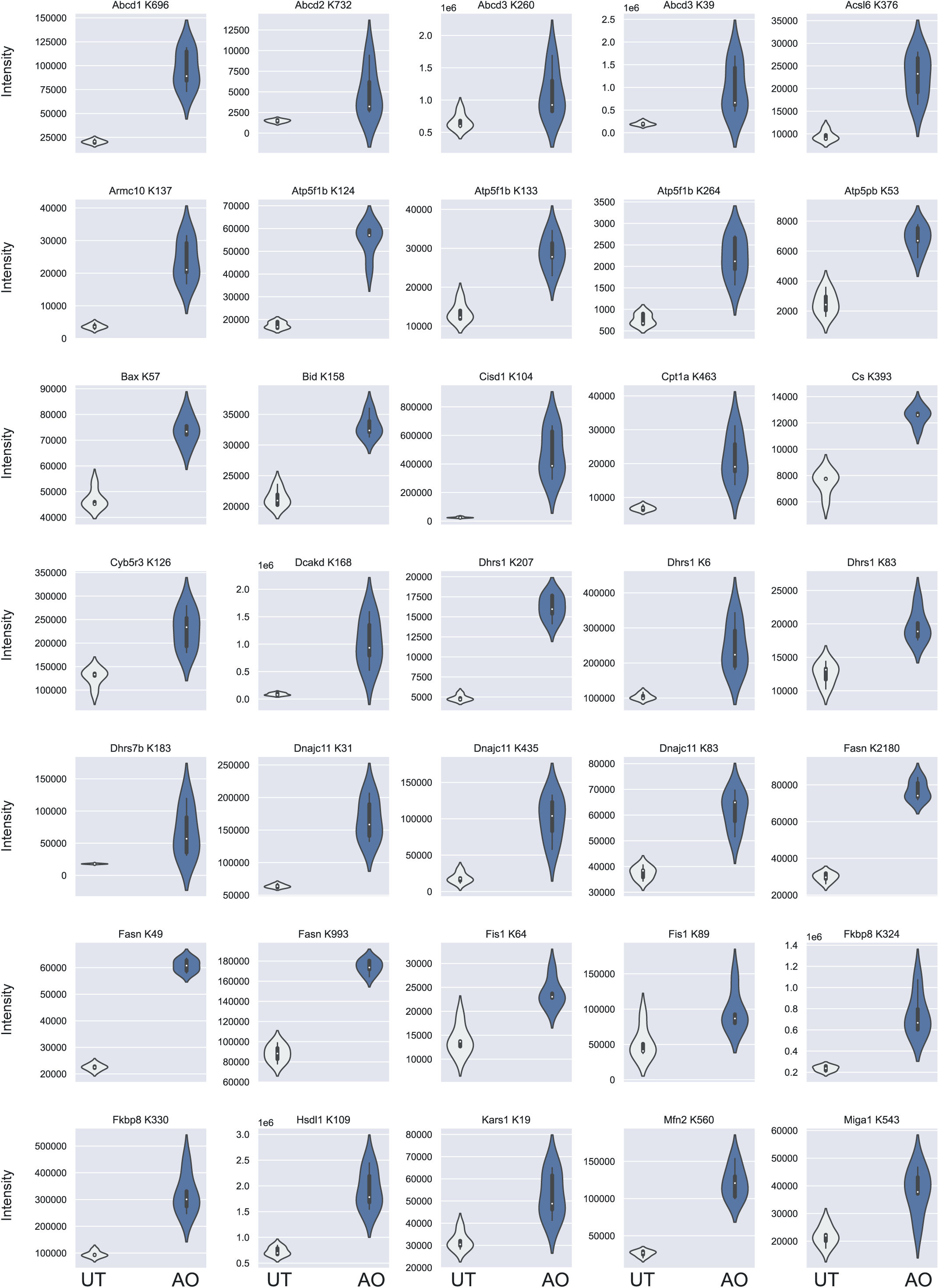
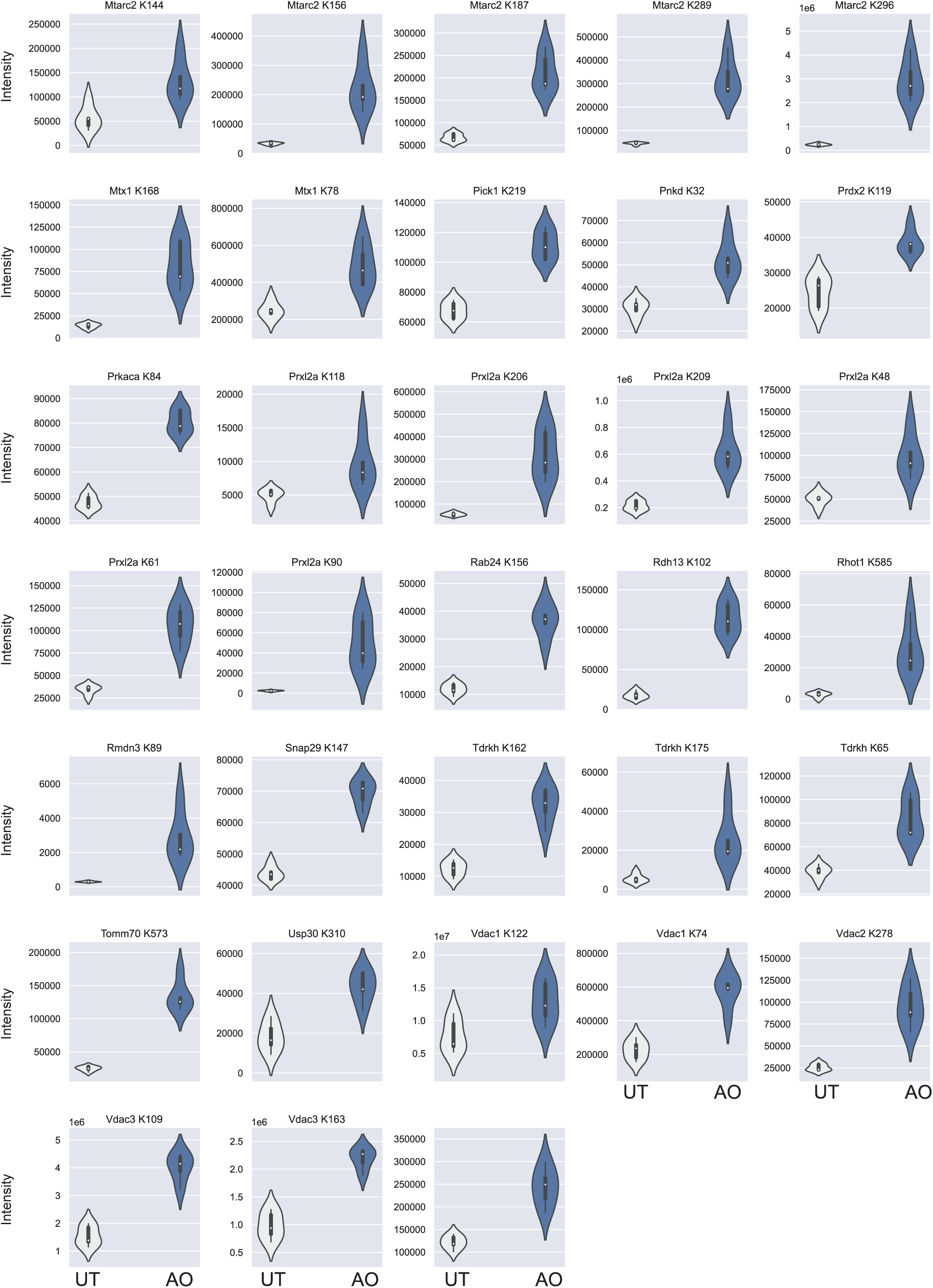
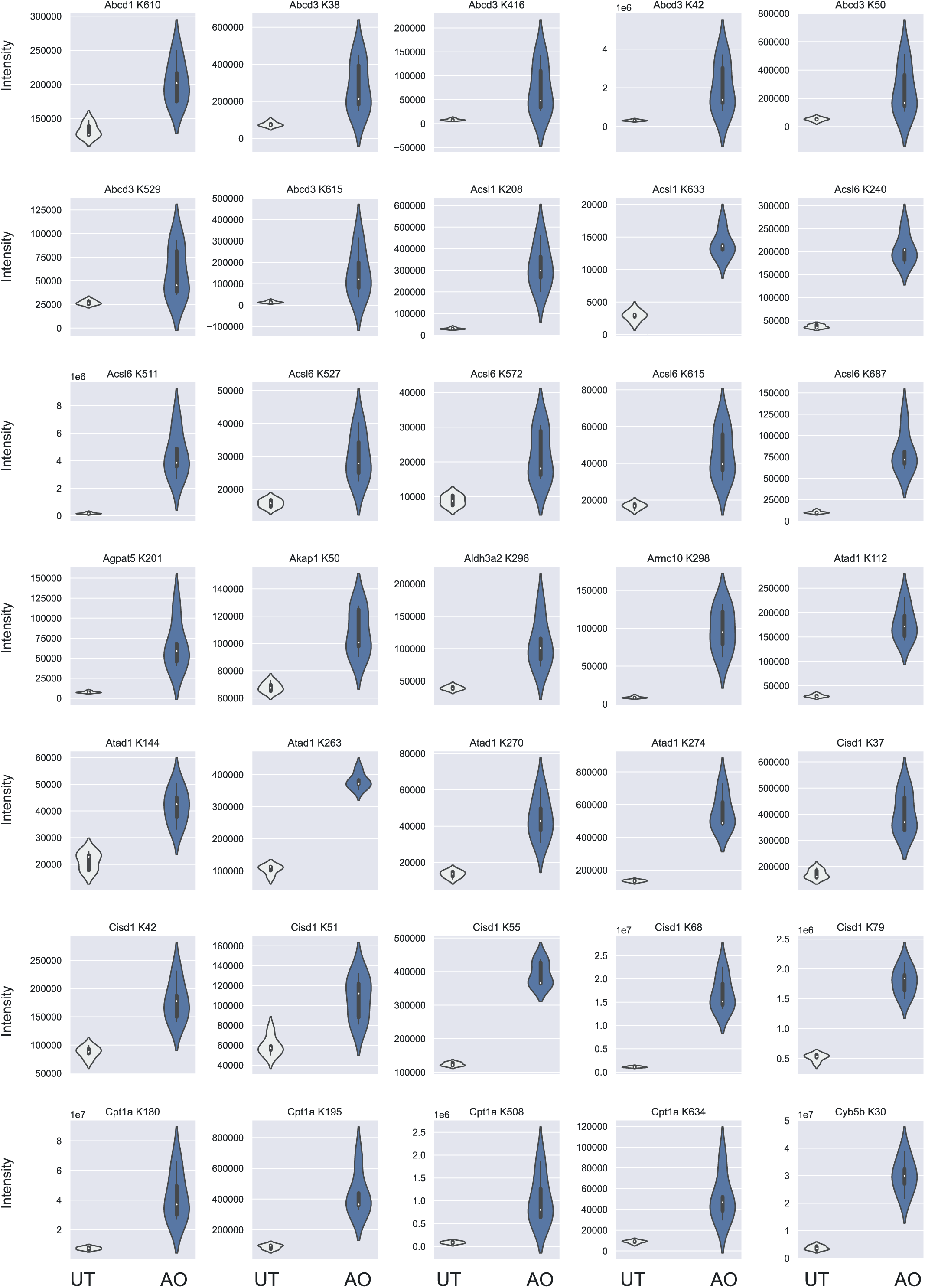
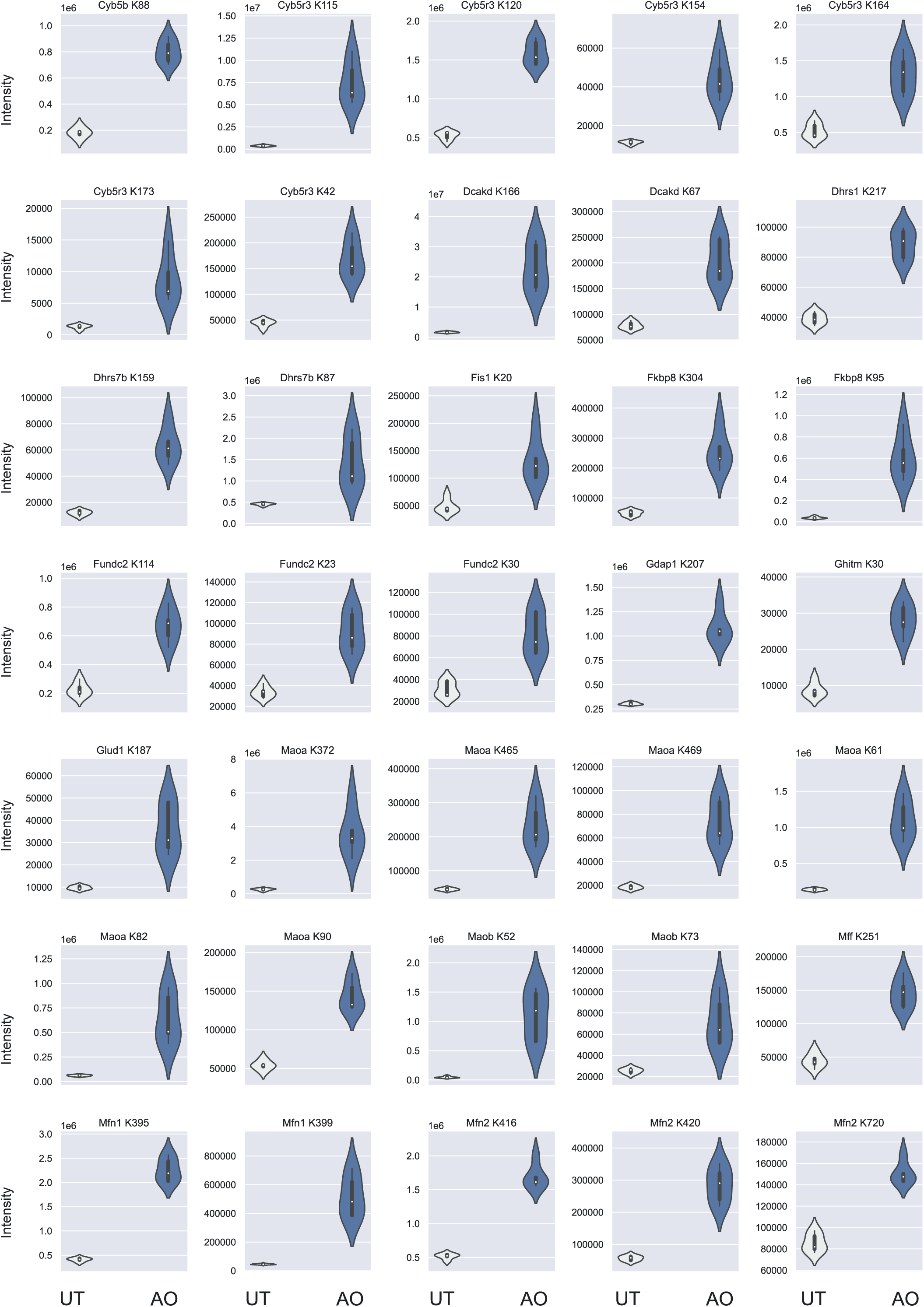
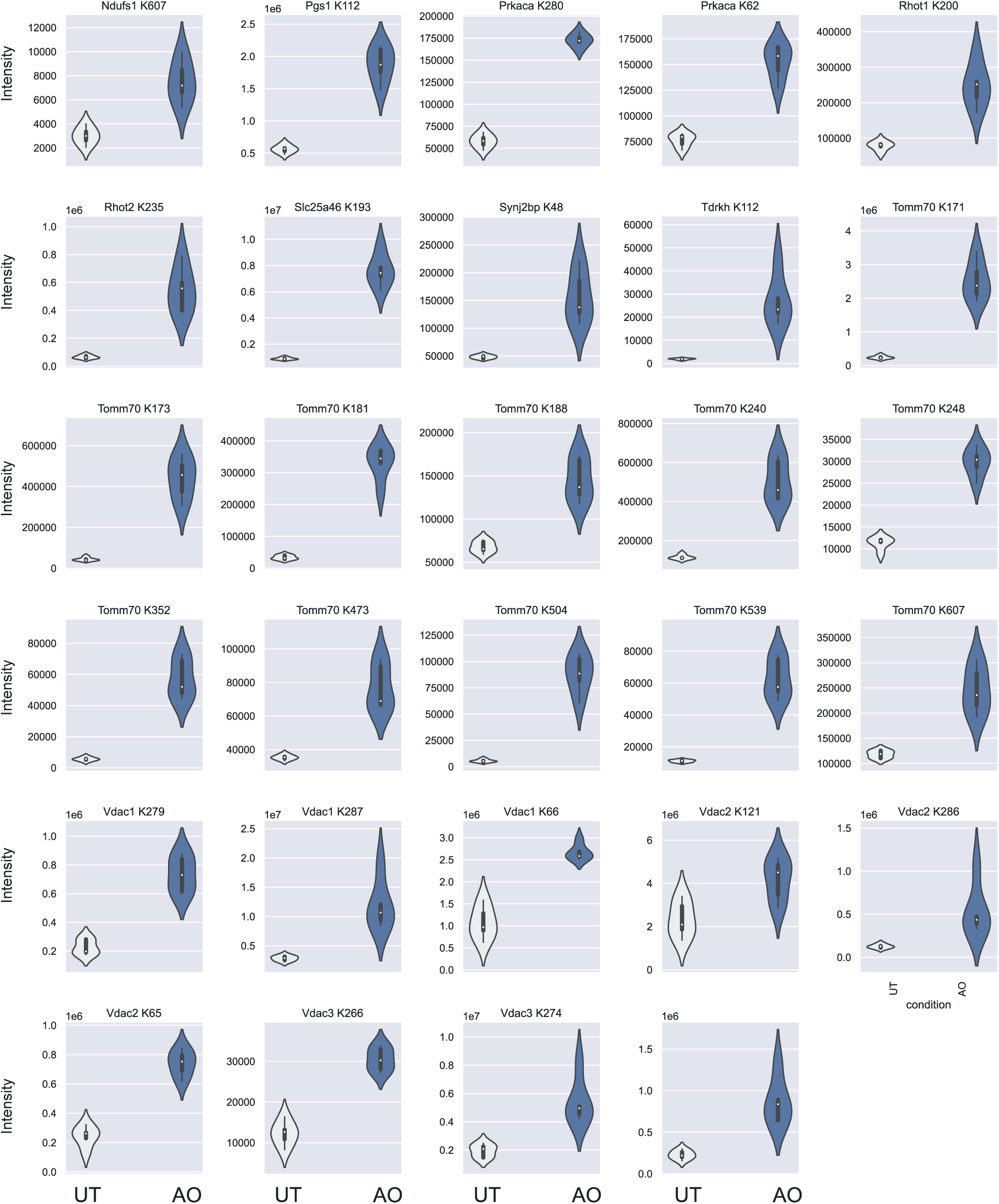
New and previously annotated Parkin mediated ubiquitylation in neurons. Grid of violin plots of significantly changed proteins with log2 fold change greater than 1.0 derived from the experiments described in Figure 8A. (A & B) Novel sites highlighted by the CURTAIN-PTM analysis. (C to E). Previously highlighted sites (25).

## Supplemental dataset legends

Dataset S1:

Tabulated dataset containing description and parameters for files input of (PPM1H) data for CURTAIN and CURTAIN-PTM. The parameter name as they appear on the web interface is in the “Parameters” column while further description of these parameters can be found in the “Description” column. The column names that have been selected as input are found in the “Column names”.

Dataset S2:

Tabulated text file containing all currently available data filter lists that can be used for batch selection within CURTAIN and CURTAIN-PTM. The title of the list is located in the “name” column. The biomolecular group/association category the data belong to is located in the “category” column. The individual protein or gene name in the list can be found in the “data” column delimited within the column by a “;”.

Dataset S3:

Tabulated dataset containing all the modification types (“Modification Type” column) and the name of the databases (“Database Name” column) the modifications.

Dataset S4:

Tabulated dataset containing the data used as input for CURTAIN and CURTAIN-PTM analysis of PPM1H-BromTAG experiment. The differential analysis for CURTAIN could be found in the “PPM1H-BromTAG_Total proteome” sheet while the data for CURTAIN-PTM could be found in the “PPM1H-BromTAG_Phosphoproteome” sheet.

Dataset S5:

Tabulated dataset containing the data used as input for CURTAIN-PTM analysis of the ubiquitome of WT-PINK1 and PINK1-KO mouse primary cortical neurons that are treated ± Antimycin-A/Oligomycin experiment. The data for analysis of WT-PINK1 mouse primary cortical neurons that are treated ± Antimycin-A/Oligomycin (AO) could be found in the “tTest_WT- AO_UT”sheet while the data for analysis of WT-PINK1 and PINK1-KO mouse primary cortical neurons that are treated ± Antimycin-A/Oligomycin (AO)could be found in the “tTest_WT- KO_AO_UT” sheet.

Dataset S6.

Tabulated dataset containing the package and software dependencies used in the development of various components of CURTAIN and CURTAIN-PTM

